# Integrated single-cell analyses of affinity-tested B-cells enable the identification of a gene signature to predict antibody affinity

**DOI:** 10.1101/2025.01.15.633143

**Authors:** Michele Chirichella, Matthew Ratcliff, Shuang Gu, Ricardo Miragaia, Massimo Sammito, Valentina Cutano, Suzanne Cohen, Davide Angeletti, Xavier Romero-Ros, Darren J. Schofield

## Abstract

Advancements in single-cell technologies and deep sequencing have revealed the vast B-cell repertoire arising from immunisation, enhancing the number of antibodies available for testing. However, selecting the highest affinity antibodies from many sequences is not a straightforward feat, as mechanisms sustaining affinity maturation and related markers remain under-studied. Here, we generated datasets of antigen-specific B-cells after mouse immunisation as well as re-analysed public data to identify a novel transcriptomic signature, "*High Signature*" (HS), with predictive power for high-affinity antibodies. HS, derived by integrating antibody sequences, gene expression, and affinity measurements, enabled sub-nanomolar affinity antibody selection without sequence pre-analysis. Notably, HS-expressing B-cells were 2.5 times more likely to yield high-affinity antibodies than randomly picked cells. Mechanistically, we identified RUVBL2, an AAA+ ATPase, as a primary HS modulator, suggesting its involvement in affinity maturation. Furthermore, HS applied to human PBMC data enriched high-affinity antibody expression, underscoring its potential in antibody discovery.

## Main

Identifying markers that distinguish high-affinity B-cells is essential for advancing antibody discovery and therapeutic design. Single-cell technologies and deep sequencing provide access to the immune repertoire diversity, yet prioritizing antibodies to synthesise from thousands of sequences remains challenging. Understanding the mechanisms by which B-cells mature could unveil new pathways to deliver high-affinity antibodies.

Affinity maturation occurs in germinal centres (GCs), transient structures in secondary lymphoid organs^1^. Within GCs, B-cells undergo iterative rounds of somatic hypermutations (SHMs) in the dark zone (DZ) and are subsequently selected in the light zone (LZ) by T-follicular helper cells (T_FH_). Positive selection of higher-affinity B-cells depends on both T-cell help and BCR signalling_2_. Recent studies underscored the importance of BCR in priming high-affinity B-cells for selection by calcium mobilisation^3^, prevention of apoptosis and enhancement of synergy with T_FH_^4^. Other studies highlighted how high-affinity B-cells exhibit accelerated cell cycle rates in the DZ^5^, a finding further supported by a metabolic shift from hypoxia and glycolysis in low-affinity B-cells^6,7^ to oxidative phosphorylation (OXPHOS) in high-affinity B-cells^8^. Furthermore, both CD40 signalling and BCR activation synergistically induce MYC, explaining B-cell selection in CGs and their proliferative capacity^9,10^.

Despite these advancements, the molecular underpinnings of affinity maturation and selection are not completely understood, especially towards the aim of finding novel biomarkers for antibody discovery and exploring the most downstream effectors of the MYC pathway.

Previous studies on affinity maturation relied heavily on simple, mostly monoclonal models, exploiting B-cell clonal expansion of a particular clone in mice immunized with antigens conjugated to 4-Hydroxy-3-nitrophenylacetyl (NP) hapten^3,8,11,12^. In this system, affinity response is dominated by the VH186.2 clone, with precise SHMs allowing a modest shift in affinity up to ∼40 nM^11^, suggesting mechanisms related to advanced phases of affinity maturation are unexplored. Consequently, in this setting, gene expression (GEX) profiles of higher-affinity B-cells are compared to the generic background population, rather than to a distinct subpopulation of low-affinity clones^8^. As such, a unifying transcriptional signature linked to affinity maturation and valid for multiple antigens across a broad diversity of clones into the sub-nanomolar to picomolar range, has not been described yet or used to select high-affinity antibodies.

Indeed, existing methods for selecting high-affinity antibodies remain limited to sequence-based approaches^13–15^, which are highly antigen-specific, or rely on *in vitro* evolution and physical selection^16,17^, which are time-consuming and require the user to employ multiple techniques.

Here, we leveraged real-world affinity measurements to integrate transcriptomic and BCR sequences with affinity data at the single-cell level across multiple sources, and we identified a novel antigen-and clone-agnostic high-affinity signature, named *High Signature (HS),* which allowed a 2.5-fold enrichment of antibodies with sub-nanomolar affinities compared to the natural population distribution. Predictions were also validated in a public dataset of human anti-SARS-CoV-2 antibodies with binding data^18^, where HS allowed the selection of antibodies with best binding strength.

Our findings introduce a new powerful framework for prioritizing high-affinity antibody candidates with broad therapeutic and diagnostic applicability.

## Results

### Higher burden of SHMs is a poor predictor of antibody affinity

To investigate the impact of SHMs and B-cell transcriptional activity on BCR affinity, we generated five single-cell datasets with coupled gene expression (GEX) and Variable-Diversity-Joining (VDJ) profiling from murine B-cells after immunisation with three different antigens (ST2, Ovalbumin and FN14); additionally, we re-analysed a publicly available B-cell dataset from mice immunised with Haemagglutinin (HA)^19^ (Supplementary Table 1).

Our initial findings were derived from sorted splenic antigen-positive B-cells obtained from immunised mice with human ST2 protein (ST2 BC177 or “main dataset”), a well characterised antigen that elicits a robust antibody response^20,21^ (Fig. 1a, Supplementary Table 1). IgM^-^/IgG^+^/Antigen^+^ total B-cells were sorted using antigen-streptavidin or antigen-klickmer conjugated to fluorophores (Fig. 1a-b, Supplementary Table 1, Supplementary Figure 1a). Antigen-positive B-cells were processed through 10x Chromium system to obtain paired BCR and GEX RNA-sequencing data (Fig. 1c). SHM-based phylogenetic tree clustering of BCRs (MiXCR software^22^, Supplementary Table 1) was used to prioritise highly mutated antibody sequences for expression, and drive selection towards GC antibodies^23,24^ (Fig. 1c and 1h). A total of 229 different anti-ST2 IgG antibodies were synthesised, expressed, and tested for binding and affinity using Biolayer Interferometry (BLI) and ELISA, obtaining a dataset of 165 confirmed binders, corresponding to 297 single-cells due to clonal expansion (Fig. 1d, Supplementary Table 1). Anti-ST2 clones demonstrated an affinity range of over 10^5^-fold (Supplementary Table 1, Supplementary Figure 1b). We grouped antibodies in three classes based on their distribution around the mean KD value (KD = 2.53 x 10^-9^ M, Supplementary Figure 1b, Fig. 1d). “HIGH” (KD < 10^-9^ M) and “LOW” affinity antibodies (KD > 10^-8^ M) occupied the tails of the distribution, being three times less abundant than the MID affinity class (Fig. 1e). This distribution recurred with similar trends in the validation dataset (Supplementary Table 1), containing antibodies with different antigen specificities (Ovalbumin, FN14 and HA) (Fig. 1e). In contrast to reported approaches where affinity was analysed based on individual clone expansion^8,24–27^, we selected highly diverse antibodies derived from multiple VH genes (Fig. 1g, Supplementary Figure 1c). The three affinity classes were represented across the V-genes (Fig. 1g).

**Figure 1.**
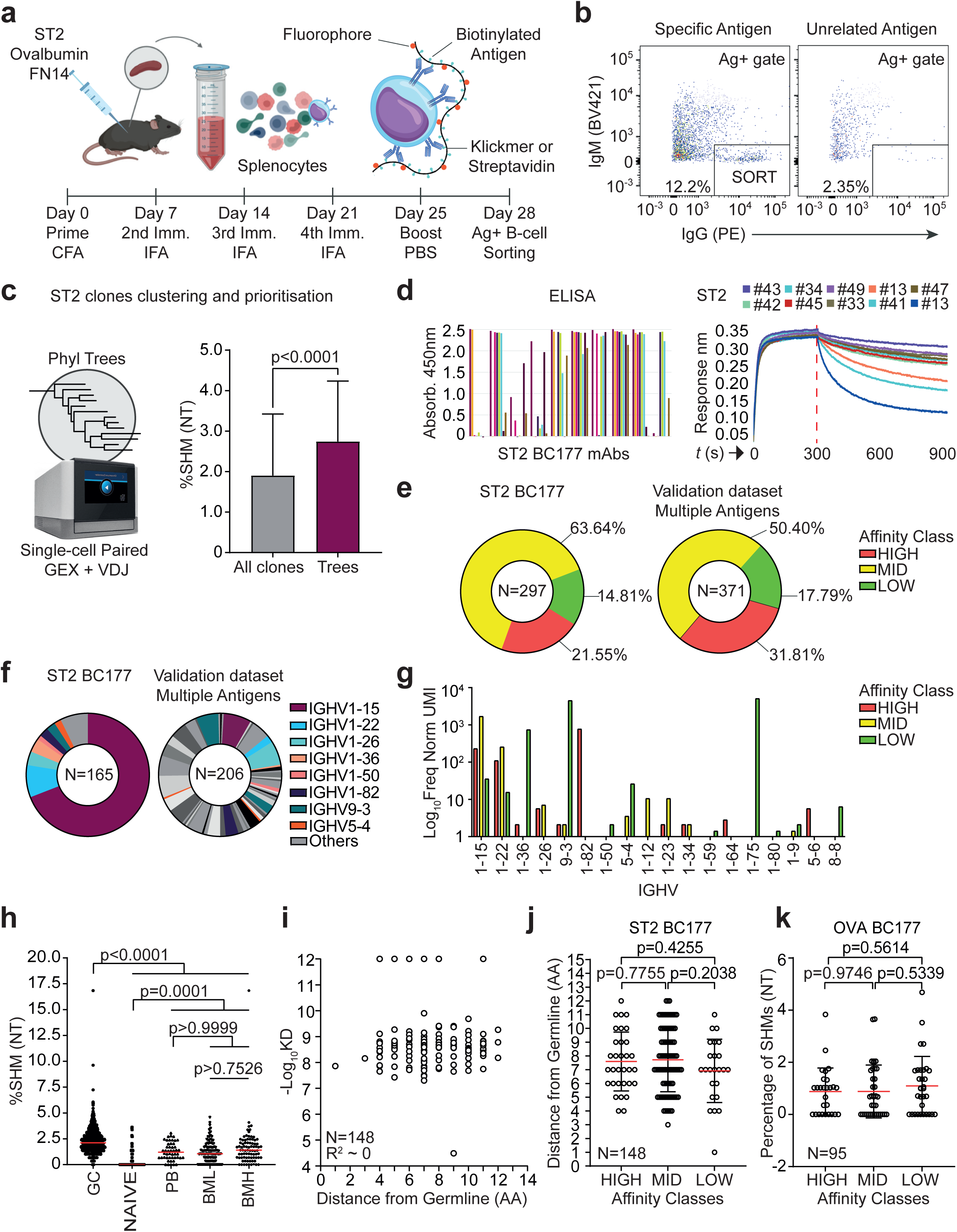
Higher burden of SHMs is a poor predictor of antibody affinity. (a) Scheme of the immunisation experiment. CD1 mice were immunised with different antigens to build datasets to discover affinity markers (ST2 BC177, N=6) or to validate findings (ST2 and OVA BC181 N=7 and N=7, FN14 N=3, HA public data N=3). (b) Sorting gate for Ag+ B-cells. Splenocytes from mouse spleens were pooled (N=2 each sample; FN14 N=1 per sample) and Ag+ cells sorted with respect to a negative control (non-specific antigen) to evaluate background. (c) Coupled analysis of single-cell RNA-seq and VDJ and subsequent phylogenetic tree clustering. Levels of SHMs are higher in phylogenetic trees with respect to the rest of the clones. N All clones = 1217, N Tree Clones = 188. Mann-Whitney U-test Two-tailed. (d) ELISA and Octet (BLI) curves from representative anti-ST2 antibody clones (BC177). Antibodies were expressed from phylogenetic trees. (e) Distribution of affinities found in B-cell clones. Affinity was divided in the following classes: HIGH = KD < 10^-9^ M, MID = 10^-9^ M < KD <10^-8^ M, LOW = KD > 10^-8^ M. (f) Pie chart illustrating observed diversity of V genes in main dataset (ST2 BC177) and validation dataset (multiple antigens). (g) Distribution and UMI frequency of V genes per affinity class. (h) Amount of SHMs found in different type of B-cells, annotated with GEX data. N GC = 753, N NAÏVE = 138, N PB = 48, N BML = 106, BMH = 82. ANOVA/Kruskal-Wallis test, Adjusted P-values. Median is shown in red. (i) Absence of correlation between number of SHMs and affinity. R squared = 0.0009624. N = 148. (j) Number of amino acid SHMs per affinity class in ST2 BC177 unique clones. Despite the presence of a trend, it was not possible to use number of SHMs (both NT and AA) to infer clones with higher affinity. Mann-Whitney U-test, two-tailed. N HIGH = 32, N MID = 92, N LOW = 24. Mean is shown in red, SD range shown in black. (k) Percentage of nucleotide SHMs in Ovalbumin unique clones. Mann-Whitney U-test, two-tailed. N HIGH= 27, N MID = 36, N LOW = 32. Mean is shown in red, SD range shown in black.

We used the BC177 dataset to test whether the general levels of SHMs correlated with affinity. Our findings confirmed significantly higher mutation rate in GC-derived BCRs as compared to other B-cell types (Fig. 1h); nevertheless, we could not observe any correlation between affinity and SHM burden within GC-derived antibodies (Fig. 1i), resulting in no significant difference between classes in both the ST2 (Fig. 1j) and the Ovalbumin (Fig. 1k) datasets. Similar findings were observed in antibodies derived from the largest ST2 phylogenetic tree #15 (Supplementary Figure 1d, Supplementary Table 1). These findings suggest that affinity is dependent on the type of mutations and VH lineage rather than on the total number of SHMs within the VDJ region ^26,27^. Indeed, analysis of SHMs shared within each class revealed differential usage of amino acid changes depending on affinity (Supplementary Figure 1e). Consistently, the number of SHMs became a slightly better predictor of affinity only when top 5 class-specific mutations were considered (Supplementary Figure 1f).

Thus, these results demonstrated that the total burden of SHMs is an unreliable proxy for affinity.

### Identification of a novel transcriptional signature linked with high-affinity BCRs

The challenge of using SHM levels to predict BCR affinity highlighted the need for alternative prediction tools. We hypothesised that increased BCR affinity might trigger selective transcriptomic changes enabling the identification of high-affinity B-cells. With this in mind, we integrated antibody affinity measurements and their sequences to annotate paired GEX data (Figure 2a-c). Seurat^28^ was used to cluster B-cells from ST2 and subclusters were annotated manually in accordance with well-defined markers ^19,29–31^ (Fig. 2a-b). The antigen-positive B-cells clustered into three main groups. The largest cluster contained GC B-cells (GCB), marked by higher levels of *Aicda/Fas*, subdivided into LZ (*Cd83^hi^*, subclusters 2-3-8) and DZ (*Cxcr4^hi^*, subclusters 0-1-7-15). The second largest cluster included memory B cells (BM) with different activation states (BMH = high-activation BM, BML = low-activation BM, subclusters 9 and 6 respectively) and naïve/unswitched cells (N, subcluster 5), known to be transcriptionally close (annotated with *Ighm, Cd80, Cd38*)^32^. Lastly, we observed a small population of Plasmablasts (PBs, *Ighg1^hi^ Sdc1^hi^*) (Fig 2a-b).

**Figure 2.**
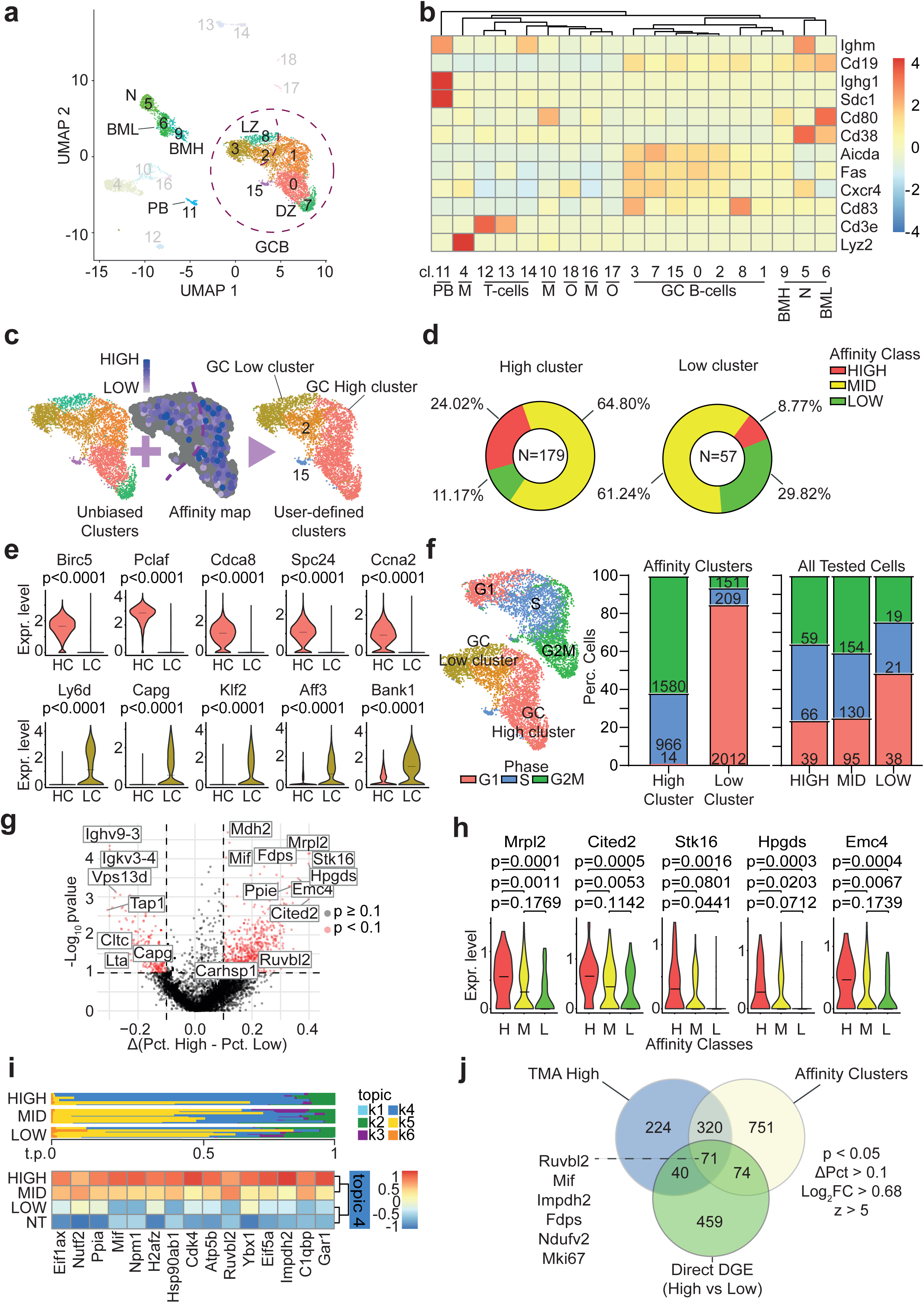
Identification of a novel transcriptional signature linked with high-affinity BCRs. (a) UMAP of ST2 BC177 data. Main clusters are identified as N = Naïve, BML = B-Memory Low Activation, BMH = B-Memory High Activation, GCB=Germinal Centre B-cells, LZ = Light Zone, DZ = Dark Zone, PB = Plasmablasts. (b) Heatmap of genes used to annotate ST2 data. Markers according to Victora et al 2010 (Cell), Victora and Dominguez-Sola et al 2012 (Blood), Mathew et al 2021 (Cell Reports). PB = Plasmablasts, M = Monocytes, O = Others, GC = Germinal Centre, BM = Memory B-cells. (c) UMAP and Affinity map of the GC from main ST2 dataset and division in affinity-based clusters (“High cluster” and “Low cluster”). Full clusters in Supplementary Figure 2a. (d) Pie chart summarising total number of tested clones and their relative distribution in the affinity clusters. (e) Violin plots showing top 5 markers identified via differential gene expression between High Cluster and Low cluster. Black horizontal bar is median. P-values from Mann-Whitney U-test, two-tailed. HC = High cluster N = 2560, LC = Low cluster N = 2372. Median is shown as a black bar. (f) Cell cycle phase distribution of ST2 dataset highlighting prevalence of high affinity cells is in the S/G2M phase. (g) Volcano plot showing markers from a new differential expression analysis between high affinity cells and low affinity cells (cells tested for affinity). Y axis = -Log_10_ (p-value), X axis = Pct.High – Pct.Low, difference in percentage of cells expressing the gene in each group. (h) Violin plots showing top 5 markers discovered with direct DGE experiment on tested cells, demonstrating high correlation and scalability with the affinity classes. Median shown. Mann-Whitney U-test, two-tailed. N HIGH = 58, N MID = 177, N LOW = 42. Median is shown as a black bar. (i) Topic model analysis to reveal additional class of markers and heatmap focussing on topic 4 showing relative expression of selected markers on the tested dataset. (j) Venn diagram showing intersection of significant markers from all analyses. Highlighted are some of the common genes that are representative of different ontology categories discussed in Figure 5. Legend describes filtering method for gene lists. P = p-value, ΔPct = difference in percentage of cells expressing a gene between group 1 and 2, FC = Average Fold Change, TMA = Topic Model Analysis, DGE = Differential Gene Expression, z = z-score.

To fully exploit our single-cell dataset and define signatures associated with affinity data, we adopted several bioinformatics strategies. Firstly, we performed differential gene expression (DGE) analysis between *High cluster* (subclusters 0-1-7) and *Low cluster* (3-5-6-8-9-11), two user-defined areas within DZ and LZ/BM/PB respectively, derived by annotating affinities (Fig. 2c, Supplementary Table 3, Supplementary Figure 2a) (here named “Affinity Clusters” method). This division generated two macro-clusters containing thousands of cells representing either low affinity (LOW) or high affinity (HIGH) classes (Fig. 2d-e), reflecting trends in affinity-tested cells (Supplementary Figure 2b, Supplementary Table 2). The affinity-based clusters unveiled net correlation between highest affinities (sub-nanomolar) and S/G2M cell cycle phase markers across all affinity-tested cells (Fig. 2e-f), including those from validation datasets (Fig. 2f), extending and generalising previous findings^5,8,33^. Unlike LOW or HIGH antibodies, MID clones were more homogeneously distributed across cell cycle phases and clusters (Supplementary Figure 2a, 2c, Fig. 2d, 2f). To find markers that were less dependent on cell cycle, we performed additional analyses using only affinity tested cells from the ST2 BC177 dataset (Fig 2g-i, Supplementary Table 2). These analyses did not consider cell type or position within clusters, but solely affinity annotations. DGE between HIGH and LOW B-cells (here termed “direct DGE” method) identified additional genes with diverse biological function and better discriminatory power (Fig. 2g-h, Supplementary Table 2). Remarkably, although DGE was conducted using HIGH vs LOW B-cells only, MID cells demonstrated intermediate expression levels, further strengthening the correlation of these genes with affinity (Fig. 2h).

Significant genes could be classified depending on expression across B-cells of different affinity. *Mrpl2, Cited2, Stk16 and Hpgds* presented a null median value for LOW B-cells and were decreasing substantially across the three classes of affinity, from HIGH to LOW, with moderate expression levels (Fig. 2h). *Ybx1* and *Hmgb2* showed overall higher expression levels, with low variance for HIGH; in contrast, *Ruvbl2* and *Mif* displayed similar expression levels in both MID and HIGH B-cells, albeit at higher level than LOW, suggesting a potential involvement in earlier phases of affinity maturation (Supplementary Figure 2d). Finally, genes such as *Lta* and *Cltc* were preferentially expressed in LOW B-cells (Supplementary Figure 2d).

Starting from the same HIGH, MID and LOW B-cells and aiming at fully dissecting the complex biological processes associated with affinity maturation, we applied topic model analysis (TMA) to our data (Fig. 2i, Supplementary Figures 2e-f, Supplementary Table 2). TMA is a machine learning method used to uncover hidden topics in text documents^34^ which recently allowed Chen and colleagues to elucidate the role of oxidative phosphorylation (OXPHOS) in B-cells bearing affinity-enhancing mutations^8^. Six topics, differentiating the groups with different power, were identified. Notably, analysis of topic k4, which showed clear expansion across HIGH B-cells, uncovered additional markers associated with high affinity, including OXPHOS-associated markers such as *Atp5b* (Fig. 2i). TMA also identified genes associated to a “LOW” topic, k5. Cells contributing to topic k5 were mainly BM, in contrast to k4 topic, that was enriched by DZ GC B cells (Supplementary Figure 2f).

Finally, we obtained lists of genes that are either exclusive for each method or common to all of them (Fig 2j, Supplementary Table 2) and used all lists to build a predictive panel for affinity.

Overall, we derived a new transcriptomic signature able to discern B-cells based on their measured affinity that we called *High Signature (HS)*. HS is the collection of all the overexpressed genes in HIGH.

### *High Signature* is antigen agnostic

To validate HS, we performed additional immunisation campaigns using different antigens: Ovalbumin (OVA) and human FN14^35^, two proteins of different size, origin, and function (validation dataset, Fig. 1e-f, Supplementary Table 1-2, Fig. 3a). Similarly to what we did for ST2 BC177, we expressed and measured affinities of antibodies from available phylogenetic trees to produce an annotated validation dataset (Supplementary Table 2). We also compared HS to *Low Signature* (LS), defined as the ensemble of all the genes that were downregulated in all the DGE and TMA analyses (Supplementary Table 2).

**Figure 3.**
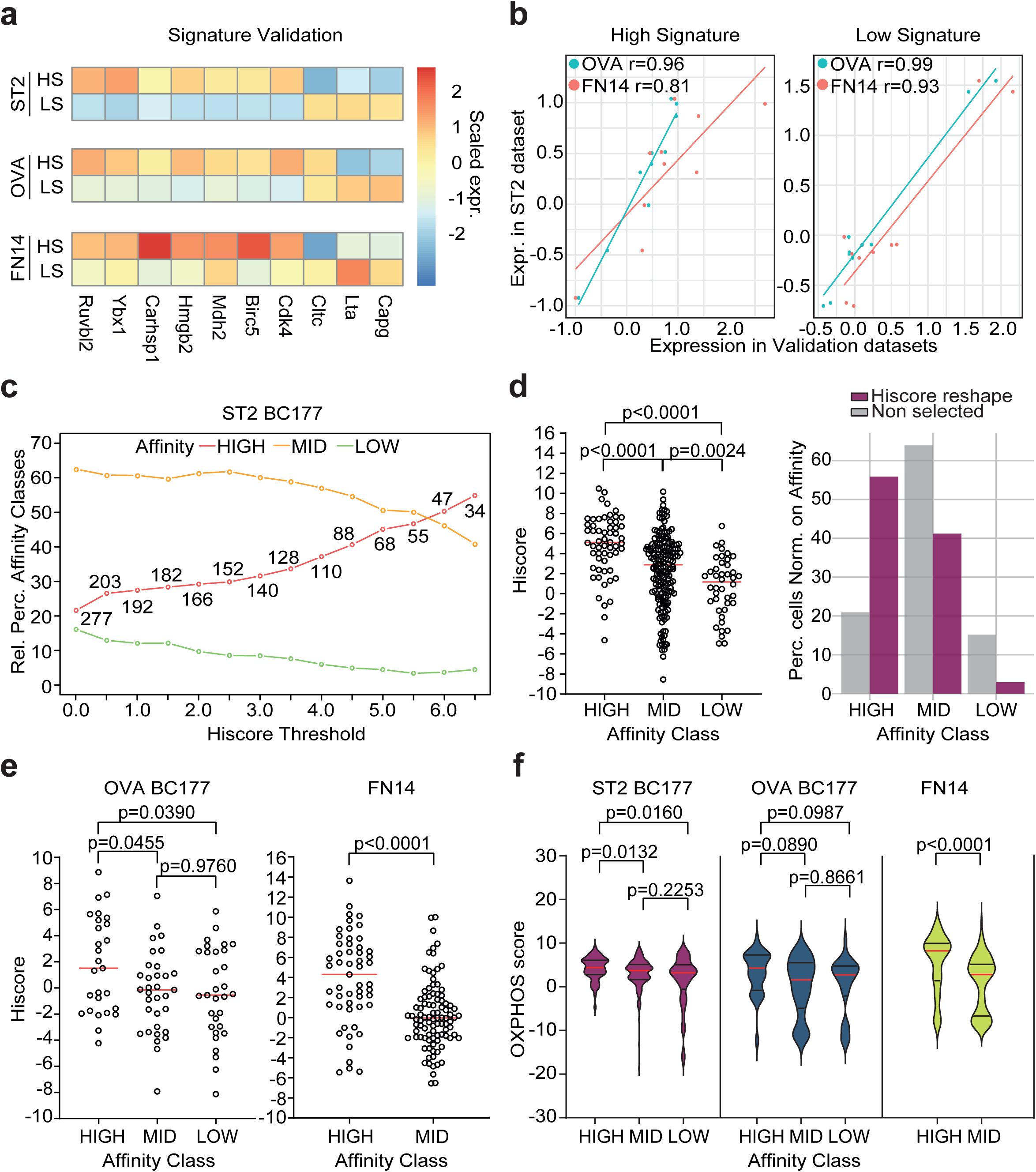
High Signature is antigen agnostic. (a) Heatmap showing relative expression of selected signature genes with high correlation in different datasets. Expression is scaled on rows. Antibodies were expressed from phylogenetic tree sequences. (b) Correlation of HS genes between ST2 and new validation datasets (Ovalbumin BC177 and FN14). r=Pearson correlation coefficient. (c) Line plot with connected dots showing how relative distribution of affinity classes is changing after applying increasing *Hiscore* (High Affinity Score) thresholds. Numbers indicate total number of cells. (d) *Hiscore* applied to ST2 and barplot demonstrating different affinity distribution before and after *Hiscore* reshape with threshold=6.5. N HIGH = 58, N MID = 177, N LOW = 42. P = p-value, Mann-Whitney U-test, two-tailed. Median is shown as a red bar. (e) *Hiscore* values on validation dataset, on cells tested for affinity. OVA: N HIGH = 27, N MID = 33, N LOW = 31. FN14 N HIGH = 53, N MID = 95. No low affinity antibodies were found for FN14 as FN14 population is made of 11 expanded clones. Mann-Whitney U-test, two-tailed. Median is shown in red. (f) Score built with OXPHOS genes from Chen et al 2021 (Nat Immunology) associated to affinity, in particular using *Cox10, Cox6b1, Ndufa4, Cox8a, Uqcrb, Atp5g2, Ndufb5, Cox6c, Atp5e, Atp5h.* Same statistical tests and number of cells of (d) and (e). The red bar represents the median, while the black bars indicate the interquartile range.

First, by setting thresholds for a few selected key genes, we identified two distinct cell sub-populations that expressed either HS or LS (Figure 3a, Supplementary Figure 3a, Methods). Genes used for HS and LS calls, such as *Ybx1* and *Lta,* were selected from all significant genes (Supplementary Table 2) and are shown in Fig. 3a-b. Selection criteria were based on GEX variance, GEX levels, or function (Methods, Supplementary Figure 3b). HS and LS genes showed strong correlation across datasets, highlighting their value as informative tools for inferring high-affinity B cells (Fig. 3b, Supplementary Figure 3c)^36^.

This property was leveraged to create *Hiscore*, a numeric score based on the sum of log-normalised, scaled GEX of HS markers, manually curated to be representative of multiple ontology categories and maximise correlation scores (Fig. 3b-c, Methods). Composite scores similar to *Hiscore* have emerged as a powerful approach to detect biological signatures across datasets and several cell types^37–39^. Increasingly high thresholds of *Hiscore* profoundly reshaped the affinity of the population towards HIGH in the main dataset (Fig. 3c-d). With respect to the original anti-ST2 population obtained by selecting clones from phylogenetic trees, the *Hiscore* population contained ∼2.5 more HIGH antibodies, and 3 times less LOW clones (Fig. 3d). Moreover, *Hiscore* could discriminate cells based on affinity also in the OVA and FN14 datasets, highlighting a significant difference between HIGH and MID-LOW B-cells (Fig. 3e).

Finally, we investigated the ability of the previously described OXPHOS process and of cell cycle to separate cells based on their affinity, as done for *Hiscore*. To this end, we built scores based on main markers^8,40^. OXPHOS genes and cell cycle phase could only partially differentiate affinity classes and only in some, but not all, datasets, confirming that our *High Signature* is based on a more comprehensive ensemble of biological pathways that can fully capture GEX changes taking place during affinity maturation (Fig. 3f, Supplementary Figure 3d).

Our data confirm HS and *Hiscore* can accurately identify high-affinity B-cells across different antigens and diverse BCRs, implying their general applicability within the RIMMS (Repetitive Immunisations at Multiple Sites) immunisation protocol^41^ (Methods) in WT mice.

### *High Signature* can predict antibody affinity independently of BCR sequence pre-clustering

Clustering based on full VDJ sequence might become challenging with larger repertoires^42^. Having carefully validated HS and *Hiscore* signatures’ ability to discriminate between antibodies of known affinity, we now wanted to assess whether we could use them to predict affinity entirely based on GEX data without previous BCR clustering. Therefore, we ran two additional immunisation campaigns (BC181, Supplementary Table 1) using both OVA and ST2 antigens. Cells were assigned to either HS or LS, resolving separate populations that could be visualised on the UMAP plot, and that correlated with *Hiscore* expression (Fig. 4a-b). Like previous datasets, HS cells showed a prevalence of S/G2M cell cycle phase, whilst LS cells were primarily in G1 phase, as expected (Fig. 4c). To validate the prediction, we expressed antibodies associated with either HS or LS call and tested them for binding and affinity (Fig. 4d, Supplementary Table 1). Results for 111 unique binders showed that 70% of antibodies with measured high affinity were correctly predicted by HS.

**Figure 4.**
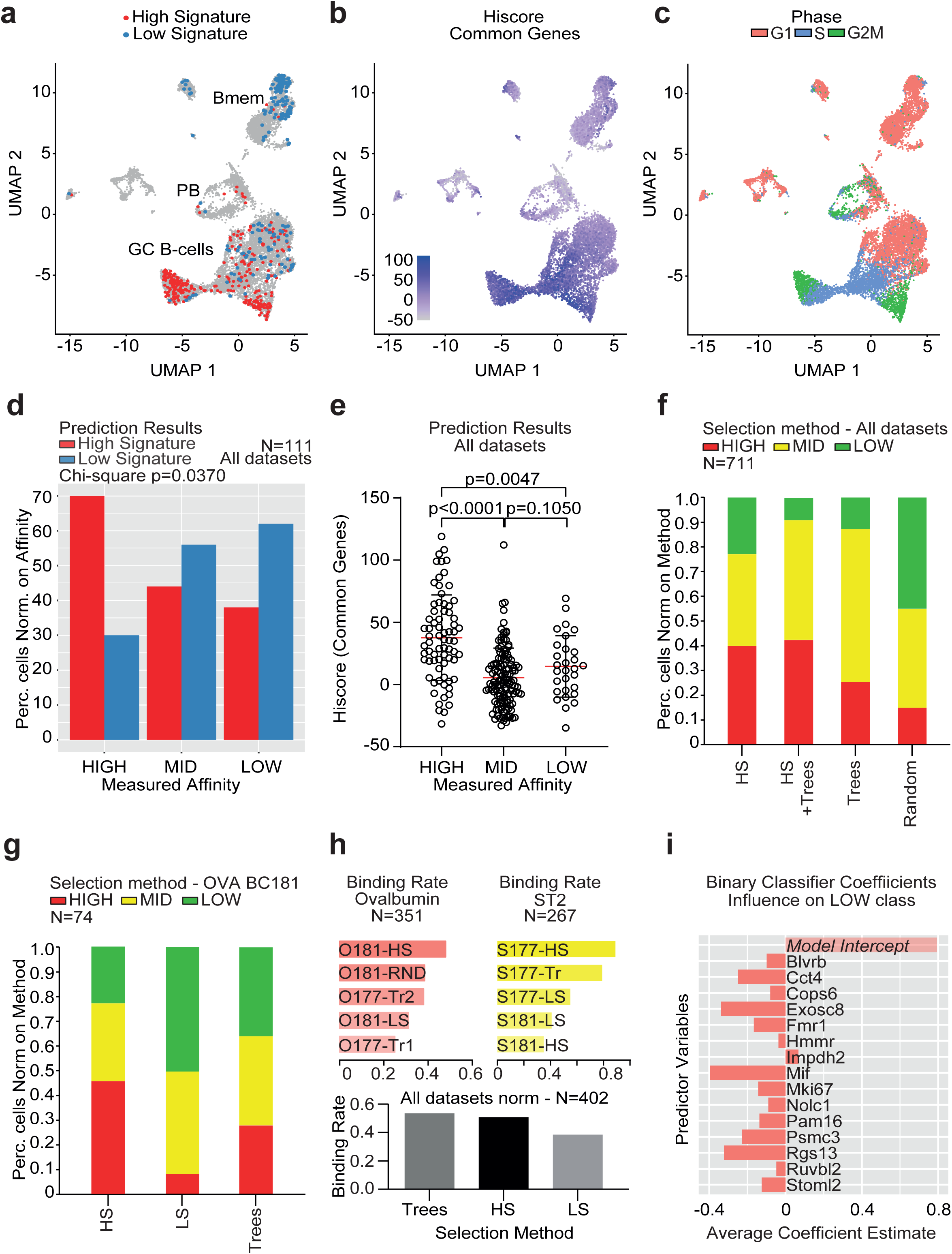
High Signature can predict antibody affinity independently of BCR sequence pre-clustering. (a) *High Signature* and *Low Signature* were used to predict cells to pick in new immunisation experiment using Ovalbumin (OVA BC181, shown in figure), ST2 (ST2 BC181 and ST2 BC177, Supplementary) and antibodies against HA (public data^19^). (b) Feature map based on *Hiscore* calculated on all common genes (Fig. 2j). (c) Phase distribution of OVA BC181 cells. (d) Aggregated results of affinity predictions for HS and LS. Data are normalised on affinity class to show relative percentage of successful predictions. N total unique antibodies = 111. N HS = 28 HIGH, 26 MID, 16 LOW. N LS = 7 HIGH, 19 MID, 15 LOW. Chi-square test of independence. (e) Highest levels of *Hiscore* in HIGH B-cells with respect to MID and LOW clones. Mann-Whitney U-test, two-tailed. N HIGH = 72, N MID = 125, N LOW = 29. Median is shown in red; SD range is represented in black. (f) Stacked barplot comparing different picking methods and subsequent shape of the affinity distribution. Statistical tests in Supplementary Figures 4a-c. (g) Stacked barplot comparing affinity distribution of OVA BC181 experiment using different methods, suggesting HS is useful in datasets exhibiting poor clustering tendency. (h) Binding rates of different batches of anti-ST2 and anti-OVA antibodies (top). Bottom: aggregated binding rate of all experiments divided by picking method. (i) Coefficients of the Multinomial Logistic Regression model. Negative values mean the gene (predictor variable) has reduced association to the reference outcome (LOW), therefore marking high affinity.

As expected, LS had similar prediction power on measured low-affinity antibodies, skewing the LOW count to ∼65%, and defining a distribution that was significantly different from HS as per Chi-square test (Fig. 4d). Both populations had comparable numbers of MID affinity (nanomolar antibodies) reflecting the nature of the most abundant affinity class (Fig. 4d). As expected, the *Hiscore* distribution was significantly different in HIGH versus MID and LOW B-cells (Fig. 4e). To compare the effectiveness of HS calls to other picking methods, we expressed additional antibodies from BC181 study by either random picking or by manually selecting based on phylogenetic trees (Fig. 4f, Supplementary Figure 4a-b). According to our hypothesis, HS pick gave the best results, that were only minimally improved when combined with phylogenetic tree clustering (Fig. 4f). In contrast, sequences picked randomly produced 3-times less HIGH affinity antibodies (Fig. 4f, Supplementary Figure 4a). OVA BC181 is a working example of how HS can be particularly useful when antibody sequences do not cluster efficiently (Fig 4g, Supplementary Figure 4c). Indeed, we observed three times less nodes per tree in OVA BC181 with respect to ST2 BC177 and still HS allowed for robust selection of high-affinity antibodies (Supplementary Figure 4d, Supplementary Table 1). Moreover, the average affinity for clones clustered in trees was lower in OVA than ST2 (Fig. 3d vs. Fig. 4g).

We also investigated the role of HS and LS in finding true binders (verified *in vitro* binders) (Fig. 4h). Indeed, data presented so far relied only on verified antibody binders, but hit rates widely varied across the different picking methods (Fig. 4h). Noticeably, HS appeared not only to be the best method to enrich for HIGH antibodies, but also for selecting true binders when compared to LS and random pick (Fig. 4h). Furthermore, antibodies selected depending on HS expression demonstrated similar binding rates to those expressed after tree clustering (Fig. 4h). This remarkable feature of HS-selected clones likely stems from their biological nature and further supports their connection to affinity. It is plausible to hypothesise that clones expressing higher levels of HS markers have experienced longer antigen exposure within the GC, thus resulting in higher affinity and consequently higher *in vitro* hit rate, when compared to other selection methods^43^.

To further improve HS prediction performance, we applied multinomial logistic regression (MLR) for binary classification (HIGH vs. LOW) on all available data (Figure 4i, Methods). The model was optimised using top differentially expressed markers from all marker discovery methods and subsets of common genes (Supplementary Figure 4e, Fig. 2j, Fig. 3a-b, Supplementary Table 2). The best predictive power was obtained from a subset of HS common genes (Fig. 2j, “Selected genes”), yielding an average AUC of 0.818 (train) and 0.730 (test), and sensitivity/specificity of 0.757/0.681 and 0.732/0.669, in train and test datasets respectively, outperforming HS+LS calls in specificity while keeping similar sensitivity (Supplementary Figure 4e-f). Probability predictions applied to MID clones were consistent with their broad position in the GEX spectrum (Supplementary Figure 4g, top panel). Overall, probability densities enabled good class discrimination, suggesting p ≥ 0.85 cutoff to be an effective method for selecting HIGH affinity antibodies in future screenings (Supplementary Figure 4g, bottom panel).

In summary, we used HS to effectively predict sub-nanomolar affinity antibodies within a B-cell population of unknown affinities.

### RUVBL2 and its downstream targets dominate High Signature in B-cells

To further our understanding of the biological and molecular pathways underpinning HS and thus high affinity, we conducted a pre-ranked Gene Set Enrichment Analysis (GSEA) ^44,45^ on HS versus LS B-cells using our main ST2 dataset (Fig. 5a-b). To provide a comprehensive view of the molecular mechanisms at play, we leveraged both the Hallmark (MH, Fig. 5a) and Targets (M3, Fig. 5b) gene sets^46^. Results demonstrated a strong involvement of MYC and E2F signalling in HS-expressing B-cells, together with active cell cycle genes (G2M Checkpoint and Mitotic Spindle terms) (Fig. 5a). In GC B-cells, MYC is a critical mediator of cell survival and cell-cycle re-entry, and it is triggered synergistically by both BCR and CD40 signalling ^47^. Within DZ B-cells, *MYC* expression is repressed by BCL6^9^. Our data suggest a role for MYC targets (downstream signalling) at later stages of affinity maturation, as HS was more highly expressed among B-cells with higher affinity. Moreover, E2F transcription factors control cell cycle progression and B-cell maturation by determining thresholds for antigen-induced T-cell proliferation^48^. Another Hallmark with high gene ratio (∼0.8) was DNA Repair (Fig. 5a), a fundamental process involved in VDJ recombination, Class-Switch Recombination and SHMs^49^. Specific factors coordinating the integration of BCR/T-cell signalling with DNA repair processes during the selection of higher affinity B-cells are currently being investigated^49–51^. Finally, with moderate gene ratios, we could confirm the previously identified OXPHOS and mTORC1 signalling pathways, which are also linked to increased mitotic activity in GCs ^8,12^ (Fig. 5a).

**Figure 5.**
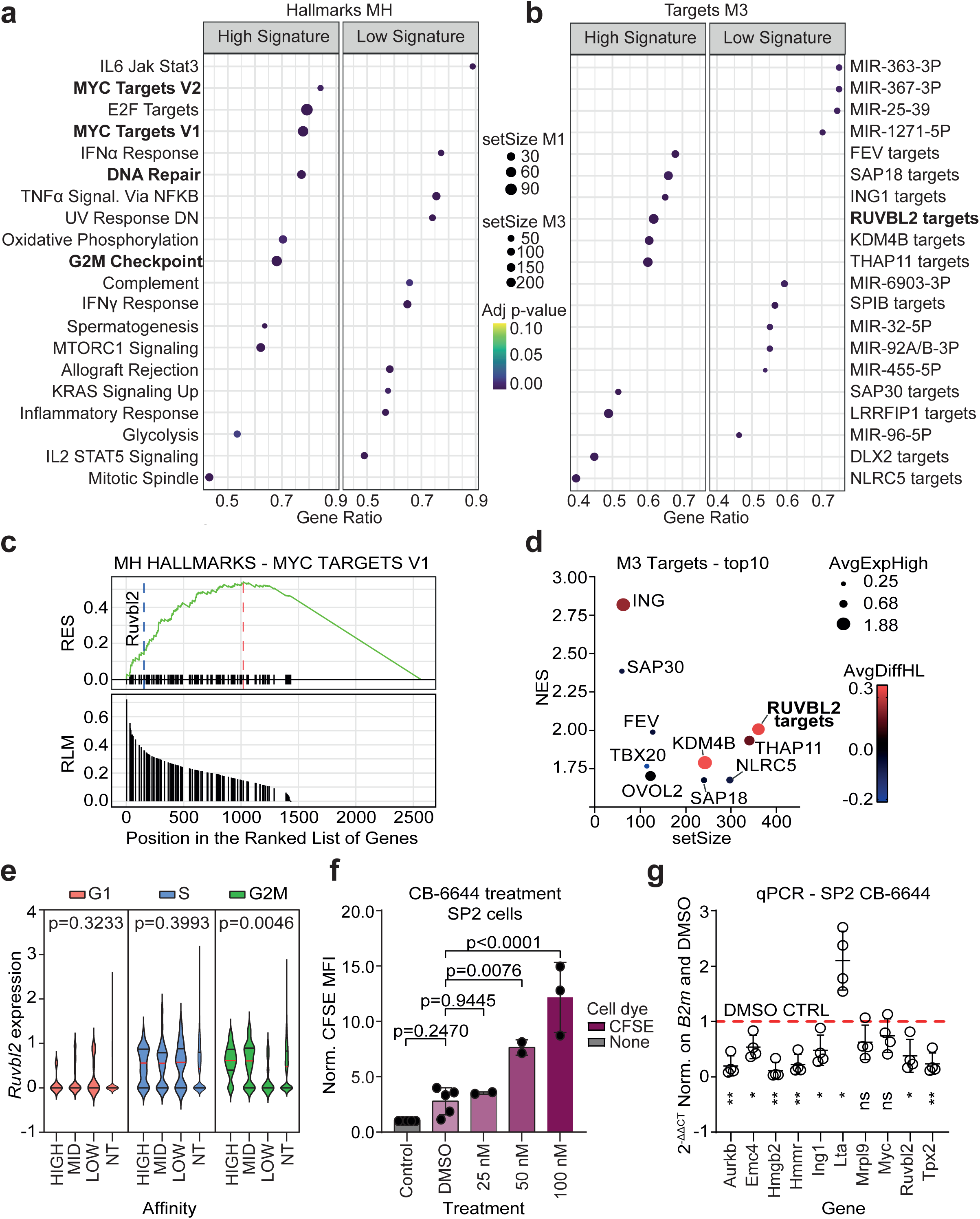
RUVBL2 and its downstream targets dominate High Signature in B-cells. (a) and (b) GSEA analyses of ST2 data, HS vs. LS, Hallmarks (MH) and Regulatory Target gene sets (M3). In bold terms that are further investigated in the manuscript and that present a biological link. (c) Running enrichment score (RES) and Ranked List Metric (RLM) from Myc Targets V1 gene set. Highlighted is *Ruvbl2* (blue dashed line) and ES (red dashed line). (d) Multidimensional dot plot illustrating top 10 terms for M3 ranked by descending Normalised Enrichment Score (NES). AvgExpHigh = Average expression level of the target genes found in the ST2 dataset; AvgDiffHL = Average differential expression of target genes between High Affinity and Low Affinity cells in ST2 dataset. All terms refer to targets of the represented genes. (e) *Ruvbl2* expression grouped by Phase and Affinity in ST2 dataset. NT = Non tested. Kruskal-Wallis test. Multiple comparisons with FDR, G2M = HIGH vs LOW q-value 0.0062, MID vs LOW q-value = 0.0195, MID vs NT q-value = 0.0424. The rest of comparisons is non-significant (q-values > 0.05). N G1 = HIGH 7, MID 35, LOW 14, NT 3355. N S = HIGH 23, MID 60, LOW 17, NT 1788. N G2M = HIGH 28, MID 82, LOW 11, NT 1772. Median is shown in red while the interquartile range is included within the black bars. (f) Normalised Mean Fluorescence Intensity (MFI) of CFSE in SP2 B-cell line treated with either CB-6644 Ruvbl2 inhibitor or DMSO. The higher the MFI the lower the proliferation rate. Normalised on negative control. N Control (no treatment) = 5, N DMSO = 5, N CB-6644 25 nM = 2, N CB-6644 50 nM = 2, N CB-6644 100 nM = 3. ANOVA one-way, Dunnett’s test, multiple comparisons. Mean ± SD is shown. (g) RT-qPCRs of selected HS genes in SP2 cells (mouse B myeloma from spleen) treated with 250 nM of CB-6644 RUVBL2 inhibitor for 48h. N = 4. Paired t-test, two-tailed. Normalised on *B2m* and DMSO. Mean ± SD is shown.

By analysing Targets M3, upregulated in HS, we identified an enrichment in RUVBL2 targets (Fig. 5b, Supplementary Table 2). Indeed, within the top10 protein effectors ranked by Normalised Enrichment Score (NES), RUVBL2 was the only known MYC target (Fig. 5c-d) ^52–54^. Although the link between MYC and RUVBL2 was only studied in other systems,^14,51,52^ RUVBL2 targets had high basal expression levels in HIGH B-cells, and their expression was significantly higher in HIGH *vs* LOW cells (Fig. 5d, Supplementary Table 2, Fig. 1j). RUVBL2 (also known as Reptin, TIP48, TIP49B, INO80J in humans) is a member of the conserved AAA+ family (ATPases Associated with diverse cellular Activities) and plays a role in various cellular processes such as DNA Repair^55^, transcriptional regulation^56^ and oncogenic transformation^52,57^, having therefore the potential to link Myc signalling, cell cycle and DNA repair in HS-expressing cells. Indeed, *Ruvbl2* expression was significantly higher in mitotically active high-affinity B-cells, but almost nil in G1 phase cells regardless of their affinity (Fig. 5e). To demonstrate the direct involvement of RUVBL2 in cell cycle regulation we used SP2, a B-myeloma cell line derived from mouse spleen. By increasing the doses of CB-6644, a selective inhibitor of the active form of RUVBL1/2 (hexameric complex)^58^, the cell duplication rate progressively slowed down, as measured by retention of Carboxyfluorescein succinimidyl ester (CFSE) dye (Fig. 5f, Supplementary Figure 5a).

To demonstrate the active role of *Ruvbl2* in regulation of HS genes, we performed qPCRs on SP2 B-cell line treated with CB-6644 (Supplementary Table 2, Fig. 5g). We analysed HS genes listed as RUVBL2 targets within Target M3 and other unrelated genes used for affinity selection, and demonstrated a clear downregulation for all of them, or upregulation where appropriate (e.g. *Lta*), upon RUBVL2 inactivation (Fig. 5g). Given the high sequence identity between *Ruvbl2* in mouse and human (Supplementary Figure 5b), we sought to verify whether the same co-regulation was also conserved in human B-cells. QPCR analyses in RAMOS cells confirmed our findings, revealing additional targets (Supplementary Figure 5c). Notably, genes downregulated in HS, such as *LTA* and *ITPR3,* increased their expression when RUVBL2 was inhibited, as expected. Moreover, *MYC* and *RUVBL2* resulted downregulated as reported by previous work^56^.

Altogether, our data establish a new axis within the *MYC-RUVBL2* signalling pathway in mouse and human B-cells, suggesting RUVBL2 network as crucial for the positive selection of higher affinity B-cells.

### RUVBL2-positive cells can be used to identify HS in Human Peripheral B-cells of SARS-CoV-2 patients

Given the observed co-regulation of HS genes in human cells upon RUVBL2 inhibition, we sought to establish whether HS could be predictive of antibody affinity in human B-cells as well. Thus, we used a publicly available dataset studying B-cells dynamics in COVID-19 patients, where authors also sorted antigen-positive B-cells from PBMCs^18^. B-cells were sorted using barcoded, streptavidin-labelled Spike, NP and ORF8 of SARS-CoV-2 and other endemic human coronaviruses^18^. Data were divided according to sampling time after infection: Severe Acute (SA), Convalescent Visit 1 (V1) and Convalescent Visit 2 (V2) (Fig. 6a). As reported by the original authors, most of the observed cells were Plasmablasts (PB), B-Memory cells (BM) or unswitched B-cells (Fig. 6a). Furthermore, the SA response to SARS-CoV-2 was mainly driven by PBs, while BM dominated later stages^18^ (Fig. 6a and Fig. 6b).

**Figure 6.**
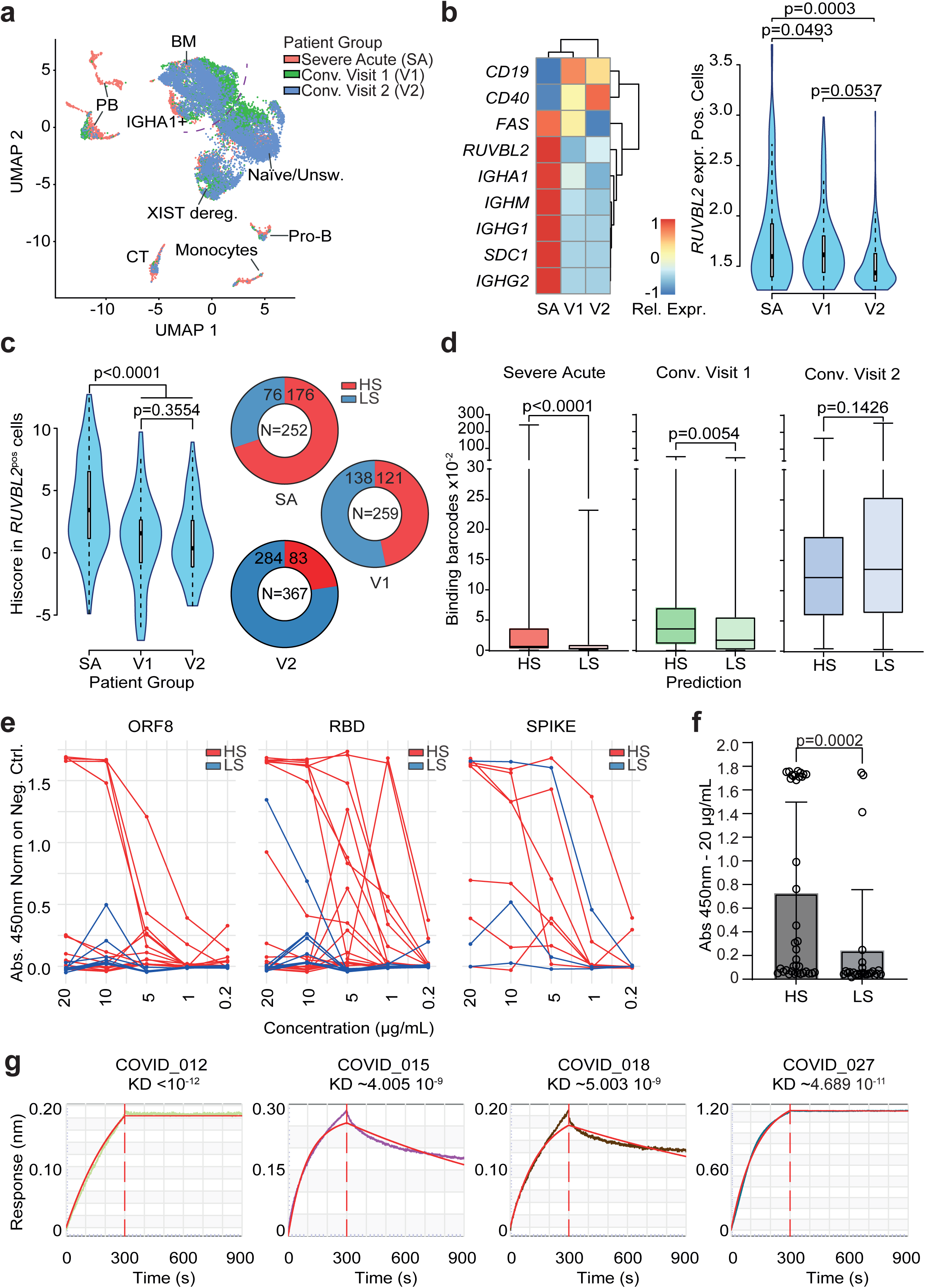
RUVBL2-positive cells can be used to identify HS in Human Peripheral B-cells of SARS-CoV-2 patients. (a) Public data from Dugan, Stamper, Li et al 2021 (Immunity) were reanalysed starting from raw data. UMAP including 4 samples per patient group annotated according to makers from the same paper and from literature. (b) Relative gene expression (scaled on samples) of genes used to annotate populations and genes such as *RUVBL2* (heatmap, left). Right: Violin plots showing expression of *RUVBL2* in *RUVBL2* positive-cells and its distribution in patient groups. Median and interquartile ranges are shown. Mann-Whitney U-test, two-tailed. N SA = 70, N V1 = 88, N V2 = 91. Median with interquartile range is shown. (c) *Hiscore* and HS applied to *RUVBL2*-positive cells across patient groups. In each group, cells were called with either HS or LS and numbers are reported in the pie charts. Mann-Whitney U-test, two-tailed. N SA = 234, N V1 = 53, N V2 = 110. Median with interquartile range is shown. (d) Antigen-binding information from the same dataset was integrated with BCR and GEX data. Box and Whisker plots of the number of total barcoded coronavirus antigens retained by BCR binding in each patient group. Mann-Whitney U-test, two-tailed. N SA = HS 146, LS = 76. N V1 = HS 121, LS 138. N V2 = HS 83, LS 284. The box represents the interquartile range, with the line inside indicating the median. The whiskers extend from the minimum to the maximum values of the dataset. (e) Antibodies called with HS and LS were expressed and tested by ELISA against corresponding coronavirus antigens (ORF8, RBD or Spike from COVID19) producing titration curves. Data are normalised on negative control (Absorbance 450nm subtraction). (f) Average absorbance value of antibodies tested in an independent ELISA experiment. Antibody concentration was 20 μg/mL while antigen coating was done at 2 μg/mL. Mann-Whitney U-test, two-tailed. N HS = 37, N LS = 26. Mean with SD is shown. (g) Octet binding profile with related KD estimate of some of the tested HS antibodies.

First, we confirmed that HS signature was also conserved in human cells, by visualizing GEX of common signature genes (Supplementary Figure 6a, Fig. 2j). Importantly, *RUVBL2* expression was significantly higher in the SA group with respect to V1 and V2 groups (Fig. 6b). This observation agrees with SA cells being assessed during infection, marking a population of cells closer to those in GCs. Unsurprisingly, the general gene landscape observed in peripheral B-cells differed from those found in GCs, with no segregation based on cell cycle phase and no expression of canonical GC markers, such as *AICDA* and *FAS* (Supplementary Figure 6b). Still, by first selecting RUVBL2-expressing B-cells, we were able to clearly identify a significant difference in *Hiscore* across patient groups (Fig. 6c). Resulting HS sub-populations were more abundant in the SA/PB group (Fig. 6c, Supplementary Figure 6c), concordant with the hypothesis that high-affinity GC cells tend to differentiate to PCs^59–61^. LS, instead, was expanded in V2 group, a subset mainly dominated by BM (Fig. 6c).

To assess whether HS and LS identified different antibody populations also in human PBMCs, we first integrated barcode-probe binding information available in this dataset with GEX and VDJ data and plotted antigen-barcodes/cell *vs* HS or LS signatures (Fig. 6d). Results showed how most binders were found within the HS-expressing cells in the SA and V1 groups, measured by a significant difference in Binding Score compared to LS (Fig. 6d, Supplementary Figure 6d). V2 group showed higher median values for both groups, and a trend in favour of LS-expressing cells as expected, but no significant difference (Fig. 6d). These data suggest HS and LS preferentially discern affinity in B-cells recently derived from GCs and not differentiated in BM. To confirm that HS was going to be a universally applicable signature that could facilitate the selection of human antibodies, we expressed and tested antibodies from both HS and LS populations for *in vitro* binding. Using ELISA against all possible antigens used for sorting^18^ we confirmed that antibodies derived from HS-expressing B-cells bound more strongly to their targets. Importantly, the number of true binders was significantly higher with respect to LS-derived antibodies, with ∼57% of the antibodies from HS-expressing cells binding to one of the antigens tested *vs ∼*15% of LS-expressing, on average, in two independent experiments (Fig. 6e-f, Supplementary Figure 6e, Supplementary Table 2). To conclude, we measured affinity for best HS-derived mAbs, observing KDs in the MID-HIGH range, further strengthening the validity of HS in human PBMCs (Fig. 6g).

Overall, our data demonstrated that HS translated from mouse secondary lymphoid organs to human PBMCs, and antibody selection based on HS significantly increased the probability of expressing high-affinity antibodies from peripheral B-cells.

## Discussion

Our integrated analyses of single-cell GEX data with BCR sequences and their affinities allowed the identification of HS, a novel transcriptomic signature to enrich for B-cells with sub-nanomolar affinity while preserving natural diversity. HS is antigen agnostic and enables antibody class affinity predictions independently of BCR sequencing pre-clustering. Our work has a clear impact for antibody discovery pipelines, especially for those not relying on further *in vitro* affinity evolution, as it directly enables a diverse repertoire to maximise chances of effective target binding and neutralization.

The affinity classification we used in our work is unique with respect to previous approaches, and surpasses previous monoclonal models used to study affinity^3,8,62^. Here we compared sub-nanomolar affinity clones (HIGH) with lower affinity B-cells that are already in the maturation process (LOW) and not to naïve cells. This allowed us to discover markers that are already active in the earliest (or active only in the latest) stages of affinity maturation, as exemplified from bottom panel of Fig. 2i when compared to all other cells (non-tested or NT, background).

The robustness and relevance of HS was demonstrated by using multiple in-house and public datasets^18,19^, including self and foreign antigens, overall achieving better performance than BCR-based sequence clustering. Selecting antibodies for expression based on sequence similarity is constrained by datasets that lack sufficient similar sequences; moreover, this selection limits the diversity of antibodies picked^15,63^. In contrast, HS leverages GEX markers representing biological processes linked to affinity maturation. This makes HS a more reliable tool to accelerate optimal selection of desired features, even when low affinity antibodies are of interest ^64^. We also foresee the possibility to use mouse reporter genes models that will allow physical sorting of high-affinity B-cells using fluorescent proteins.

Depending on the selection method and features used, HS provided up to 80% sensitivity and 70% specificity in HIGH vs. LOW predictions, without considering MID clones, for which predictions are not needed. Additional work is needed to enhance performance by identifying HS genes that most strongly drive class separation. This goal will be addressed by investigating the molecular mechanisms underlying HS and by expanding the single-cell affinity dataset to enable feature selection using machine learning techniques, including random forest classifiers, Lasso regression, and neural networks. Additionally, future analyses may help identify markers that better distinguish intermediate affinities.

We showed that SHM load is a poor way of selecting high-affinity antibodies. A significant discrimination of affinity classes was observed only when selecting the most frequent SHM changes between HIGH and LOW, and within a single phylogenetic line. Our data support the idea that only specific SHMs are contributing to affinity despite being selected, a concept still under investigation in literature^26,27,65,66^. The reason behind this behaviour might lie in neutral and bystander SHMs, which do not affect B-cell survival or antibody stability^66,67^. Therefore, an analysis of SHM frequency between affinity classes is likely to highlight what are the important mutations contributing to affinity, a process that however relies on real measurements and that is antigen specific. Thus, we propose that SHM analysis could complement selections based on HS+LS, if needed, as our signatures would identify high and low affinity B-cell populations within phylogenetic trees to be further analysed for SHM frequency.

HS is enriched in targets of RUVBL2, a highly conserved protein with a versatile role in cells and downstream target of MYC^53,68^. By selective inhibition of RUVBL2 protein we proved co-regulation of HS genes in mouse and human B-cell lines, thereby suggesting a direct involvement of RUVBL2 targets in B-cell affinity maturation via HS. Although *Ruvbl2* mutations are known to delay antibody class switching primarily due to compromised T-cell help^69^, our findings are the first to suggest a B-cell intrinsic role for RUVBL2 in affinity maturation. Our data provide novel molecular targets in the MYC-RUVBL2 axis, which we will leverage by deeper investigation *in vivo*.

Many annotated RUVBL2 targets with different functions are enriched in HS, including members of the ATP-synthase family like *Atp5a1, Atp5j2 and Atp5c1*, involved in the OXPHOS metabolic shift necessary in higher-affinity B-cells^8^. HS confirms and expands previous work, suggesting RUVBL2 might be a key player in elucidating the dynamics between the cell cycle and OXPHOS, and unravels further mechanisms overarching the positive selection of higher-affinity B-cells. Notably, the diverse ontology categories represented by HS reflect an intricate network of interconnected pathways. For example, HS includes target genes of the KDM histone demethylases family, epigenetic modulators recently linked to affinity^7^ (Figure 5d). These authors demonstrated that *Kdm2a* repression by miR-155 enhances mitochondrial fitness in high-affinity GC B-cells, further supporting connection among HS, RUVBL2, and OXPHOS^707172^

Remarkably, HS translated to human peripheral B-cells, demonstrating broad applicability to therapeutic antibody discovery and human pathologies. As HS connects high-affinity BCRs with their B-cell GEX profiles, it will allow to explore the contribution of the most reactive clones in the context of tumour-infiltrating B-cells or GC-dependent autoimmune diseases like Lupus^70^. Determining the correlation of HS with clinical scores and inflammatory signatures of such diseases will be key to identify novel mechanisms and reactive populations^71^, and to demonstrate the role of autoantibodies in diseases such as Multiple Sclerosis^72^.

Overall, HS enhances our arsenal of tools for advancing antibody discovery with therapeutic purposes and provides insights into the novel biology governing B-cell affinity maturation, which may have a profound impact on autoimmune B-cell-driven diseases, oncology, and the development of efficacious vaccines.

## Data Availability Statement

Raw data from single-cell experiments have been deposited to NCBI SRA and will be available once the manuscript is accepted for publication (BioProject PRJNA1190485). Information and data to reproduce the analyses are available in the Online Methods and Supplementary Tables.

## Code Availability Statement

Custom code used for this manuscript will be available upon request to corresponding authors. Information to reproduce the analyses has been already made available in the Online Methods section.

## Materials

Materials are described alongside relevant methods, but an additional Excel sheet summarising all Materials used with catalogue numbers is available to facilitate the reader (see **Materials Used in This Study**).

## Online Methods

### Mouse immunisations and sorting of antigen-positive B-cells

In vivo procedures performed in the United Kingdom were conducted under the authority of a Home Office issued Project License in accordance with the Animals [Scientific Procedures] Act 1986 with appropriate ethical approval. Female WT CD1 mice were immunised using RIMMS protocol (ref ^41^). Five immunisations were administered in 28 days, using 10 μg of antigen per dose according to schedule in Fig. 1a. On day 28, spleens and lymph nodes were taken, and dissected using gentleMACS (Miltenyi Biotec, Cat. No. 130-193-235). Samples used contained 2 pooled spleens each if not otherwise indicated in the text. Splenocyte underwent red blood cell lysis (Myltenyi Biotec Cat. No. 130-094-183) or Ficoll separation (see below). Cells were immediately used for antigen-positive (Ag⁺) B-cell staining and sorting. Streptavidin and antigen were assembled for at least 30 min RT at a 1:4 molar ratio with a final Streptavidin concentration of 200 nM. Cells (∼40–80 million from one mouse spleen) were stained in FACS buffer (PBS + 1% BSA). After a 30-minute incubation on ice with Jackson AffiniPure F(ab’) Goat Anti-Mouse IgG Fcγ fragment-specific (PE, 1:500 Cat. No. 115-116-071), cells were washed with 15 mL FACS buffer. This was followed by a 30-minute incubation with ThermoFisher FcR blocker (anti-CD16/CD32 Cat. No. 14-0161-82) and a wash. Antigen and antigen control were loaded onto streptavidin-AF647 (BioLegend Cat. No. 405237) or Klickmer-APC (2 µL per 5 million cells, Immudex, Cat. No. DX01K-APC-200) and incubated for 30–60 minutes on ice, followed by another wash. Additional staining included Biolegend clone RMM-1 anti-IgM BV421 antibody (1:100, Cat. No. 406517) and optional ThermoFisher Live/Dead NIR780 staining (1:1000, Cat. No. L34994) for 30 minutes on ice. After two washes, cells were filtered and sorted using BD FACSAria cell sorter. FMOs and single stains were used for compensation when required. Antigen-positive B-cells were sorted into 1.5 mL tubes using 0.1% BSA/PBS buffer as landing, spun 5 min at RT and 300 x g and resuspended in PBS for 10x Genomics experiment.

### Ficoll separation of lymphocytes from mouse spleen

Mouse splenocytes were isolated from immunized mice and divided into two 50 mL Falcon tubes. Ficoll-Paque Premium (1.084 g/mL, 15 mL/tube, Merck, Cat. No. GE17-5446-02) was used for density gradient centrifugation. Splenocytes (10⁹ cells) were resuspended in a medium containing 5 mL complete medium, 5 mL PBS, and 25 mL FBS, and gently layered over Ficoll using a 25 mL Strippette at ∼0.25 mL/s. Centrifugation was performed at 805 rcf for 30 minutes at room temperature with no brake or acceleration. The leukocyte ring at the Ficoll interface was aspirated, washed with 50 mL complete medium, and centrifuged at 453 rcf for 15 minutes (room temperature, full brake). The pellet was filtered (35 µm filter) into 15 mL tubes, washed with 15 mL complete medium, and centrifuged at 300 rcf for 10 minutes (room temperature, full brake). Viability and cell count were measured before proceeding to downstream experiments.

### 10x Genomics and Illumina sequencing

Antigen-positive B-cells per sample were used as input for 10x Genomics Chromium using ChipK (Cat. No. 1000287) and 5’ Next GEM kit v2 (Cat. No. 1000263). VDJ regions were amplified using Chromium Single Cell Mouse BCR Amplification Kit (Cat. No. 1000255). Libraries were prepared using Library Construction Kit (Cat. No. 1000190). Sequencing was performed by Illumina NovaSeq6000 in paired-end 300 cycles. NovaSeq 6000 SP Reagent Kit v1.5 (300 cycles) Cat No. 20028400 or NovaSeq 6000 S1 Reagent Kit v1.5 (300 cycles) Cat. No. 20028317 were used as needed. Minimum sequencing depth was 5,000 reads per cell for VDJ libraries and 20,000 reads per cell for GEX libraries. Read length used was 151-10-10-151. Further details on materials used in the Material Excel Sheet.

### Analyses of single-cell data using Seurat

Code is available upon request. Briefly, FASTQ files from GEX libraries were analysed with Cell Ranger, then secondary analyses were performed using R, particularly using Seurat. Data were log normalised and scaled, dimensionality reduction was applied, and clusters were found using FindNeighbours/FindClusters. Cluster density was evaluated using clustree package. Clusters were annotated in the metadata of the objects by checking GEX profile of known markers. FindAllMarkers was also used to infer most represented markers per cluster. Metadata of objects were annotated with antibody VH and VL from MiXCR or Cell Ranger VDJ clonotype tables, with phylogenetic tree information, and with affinity and binding data using barcode matching between VDJ and GEX data.

### Differential Gene Expression Analyses

DGE Analysis were conducted using cluster in Seurat Objects and affinity annotations of single cells. Affinity Clusters were defined using RenameIdents as *High cluster* (subclusters 0-1-7) and *Low cluster* (3-5-6-8-9-11) using an initial cluster resolution of 0.8. DGE was performed using FindMarkers between the two groups. For direct DGE between HIGH and LOW cells, FindMarkers was used between cells annotated as high or low affinity, regardless of their cluster or cell type. In both cases, genes showing p < 0.05 and a difference between pct.1 and pct.2 of at least 0.1 were selected as significant.

### Calculation of *Hiscore*

*Hiscore* was calculated by weighted sum of scaled GEX data from relevant Seurat objects. Two versions of *Hiscore* are used in this manuscript. *Hiscore* in Figure 3 is based on genes with highest correlation, while *Hiscore* in Figure 4 is calculated on all common genes (Fig. 2j and Supplementary Table 1). In the first instance, *Hiscore* is calculated using selected_genes vector: c("Ruvbl2", "Ybx1", "Carhsp1", "Hmgb2", "Mdh2", "Birc5", "Cdk4", "Cltc", "Lta", "Capg"). This vector contains genes that are both downregulated (*Cltc, Lta, Capg*) or upregulated in HS (rest of the genes), therefore, downregulated genes were weighted with a -1 factor in order to keep *Hiscore* positively correlated to high affinity. In the human dataset, *Hiscore* was calculated using genes in the following vector: c("MRPL2", "YBX1", "CARHSP1", "MIF", "HMGB2", "MDH2", "CLTC", "LTA", "CAPG", "RUVBL2"). Downregulated genes such as *CLTC*, *LTA* and *CAPG* were weighted with a -1 factor. *Hiscore* was used as a tool to quantify the presence of HS genes in B-cells.

### Prediction of High and Low affinity cells using *High Signature* (HS) and *Low Signature* (LS)

Cells selected with HS or LS were called as follows. HS in ST2 BC177: *Cltc* ==0 & *Ybx1* >2.5 & *Ighg1* > 0 & (*Cks2* > 1 | *Cdk4* > 1 | *Mki67* > 1). *Lta* was also used in lieu of *Cltc*, and *Cdk4* was used alternatively to *Cks2*. The thresholds for Ybx1 and Cks2 were determined by analysing the average expression level of the gene in high-affinity B-cells, and were adapted to other datasets by normalising these values on the new average expression (Gene threshold / Average gene expression level * Average gene expression level of the new dataset). For instance, Ybx1 threshold in OVA BC177 was 2.5 / 7.54 * 4.88 = 1.62. LS in ST2 BC177: (*Lta* > 0 | *Capg* > 0) & *Cks2* < 1 & *Ighg1* >0. Also, in this case values of genes were adapted to average expression levels in other datasets. *Ybx1* was selected because it had the lowest variance in HIGH cells; *Cks2* or *Cdk4* or *Mki67* are needed to select cells in active proliferation; *Cltc*, *Lta*, *Capg* are genes of LS that are set to 0 when calling HS or greater than 0 when calling LS; *Ighg1* (or other relevant antibody isotype genes) are set greater than 0 to ensure selection is done on desired type of B-cells.

### GEX Thresholds used for HS and LS selections in all datasets

ST2 thresholds are described in previous paragraph. OVA BC177: HS Lta == 0, Ybx1 > 1.62, Ighg1 > 0.43, Cks2 > 0.40; LS Lta > -.59, Capg > 1.87, Ighg1 > 0.43, Cks2 < 0.04. OVA BC181 HS Ybx1 > 2.61, Cltc == 0, Ighg1 > 0.27, Cks2 > 0.61; LS Lta > 0, Capg > 0, Ighg1 > 0.27. Cks2 < 0.61. HA HS Ybx1 > 2.74, Cltc == 0, Ighg1 > 0.024, Cks2 > 0.93; LS Lta > 1.81, Capg > 0.47, Ighg1 > 0.024, Cks2 < 0.093. FN14 HS Ybx1 > 1.36, Cltc == 0, Ighg1 > 0.03, Cks2 > 0.1; LS Lta > 0.32, Cks2 < -.01, Capg > 1.66. For human data (COVID dataset), following thresholds were applied: HS (LTA == 0 & CKS2 >1 & (IGHG1>0 | IGHG2>0 | IGHA1>0)) | (YBX1>2.5 & CKS2 >0) ; LS (LTA > 0 & CKS2 < 0.5 & (IGHG1>0 | IGHG2>0 | IGHA1>0)) | (LTA > 0 & CKS2<1). To select for RUVBL2-positive cells where appropriate we selected for IGHG1 > 0 & RUVBL2 > 0 cells.

### Topic Model Analysis

Topic Model Analyses were performed in R using the fastTopics package (https://stephenslab.github.io/fastTopics/). TMA were applied to Counts data of the annotated ST2 object using fit_topic_model function with k=6. K=6 derived from a Maximum Likelihood convergence analysis. DGE analysis was performed on the model using de_analysis function, with pseudocount = 0.1. Fit was evaluated using loglik_multinom_topic_model. DGE data were visualised using volcano_plot. Topic prevalence in HIGH, MID and LOW affinity B-cells was observed using structure_plot. To visualise top 200 cells that contributed to each topic, the output of the model fit was sorted in descendent order on topic k4 or topic k5 and corresponding barcodes annotations were extracted to be visualised in Seurat DimPlot.

### Phylogenetic tree analysis

VDJ libraries were also processed using MiXCR analyze 10x-vdj-bcr-full-length command. Phylogenetic trees are SHM-based and were find using MiXCR findShmTrees. For antigen positive B-cells, parameters were adjusted to maximise clustering. findShmTrees -f -O steps[0].algorithm.commonMutationsCountForClustering=4 -O steps[0].algorithm.maxNDNDistanceForClustering=1.5 -O steps[1].maxNDNDistance=1.7 -O steps[2].maxNDNDistanceBetweenRoots=1.0. For FN14 dataset, VDJ FASTQ data were first joined together using *cat* (terminal) and then processed using changeO in Immcantation.

### Analysis of SHMs Levels

SHMs levels were obtained as a result of the MiXCR 10x-vdj-bcr-full-length command pipeline using *exportClones -vBestIdentityPercent.* SHM levels were obtained by calculating 1 -vBestIdentityPercent values. When using phylogenetic trees, SHM levels from *exportShmTrees* or *-distance germline* (*DistanceFromGermline*) was considered. Amino acid SHMs levels were calculated based on the field *aaMutationsVDJRegionBasedOnGermline.* Total mutations per clone were counted in the full VDJ region, providing the number of total amino acid SHMs with respect to the original germline sequence.

### Analysis of SHMs frequency

SHM frequencies were also obtained using MiXCR *aaMutationsVDJRegionBasedOnGermline.* This field contains the detailed amino acid mutations of each clone with respect to the germline. The strings were split using the comma separator, then tabled using *pivot_longer* in R. Annotations from affinity were added and total number of mutations per clone per affinity classes were calculated.

### Multinomial Logistic Model

All available data were used to fit a multinomial logistic model using nnet::multinom in R. Scaled GEX data from all datasets were collected together with information on affinity class using different set of genes (Fig. 4i, Supplementary Figure 4e). The aggregated expression matrix was then divided randomly into a train (70%) and test (30%) dataset using createDataPartition. HIGH and LOW classes only were considered, and the two groups were balanced using ROSE (Random Over-Sampling Examples), which was used on train dataset only, leaving the test dataset untouched. Predictions and confusion matrixes were calculated using stats::predict and caret::confusionMatrix respectively. ROC curves and AUCs were visualised using pROC package.

Coefficients were extracted using broom::tidy on the model fit, and visualised using ggplot2. Model in Fig. 4i was then applied on all data using predict, including MID affinities, to compute probabilities of distribution and ggplot + geom_density for visualisation. Data are averages of four independent experiments.

### Gene Set Enrichment Analysis (GSEA)

GSEA analyses were performed in R using the clusterProfiler::GSEA function with the following options: pAdjustMethod = "fdr", pvalueCutoff = 0.9, minGSSize = 5. List of genes and references were obtained from the Molecular Signature Database using msigdbr::msigdbr. Dotplot and gseaplot from enrichplot package were used for visualisation.

### Serum ELISA and ELISA assays

Nunc MaxiSorp plates (Invitrogen Cat. no. 44-2404-21) were coated overnight at 4°C with 2µg/µL of antigen, 50µL per well. Plates were then washed with 1x PBS, dried, and frozen upside down at -20°C for 1 week. After thawing, the plates were washed with 1x PBS. Alternatively, plates were coated O/N at 4C and then used for blocking. Blocking was performed with 200µL of 3% milk in PBS for 1.5 hours at room temperature, followed by another PBS wash. Serum samples were diluted 1:200 in PBS with 3% milk, followed by progressive dilutions of 1:1000, 1:5000, 1:50000, and 1:500000 in the blocking solution. 50µL of each dilution were added to the wells and incubated at room temperature for 1 hour. Expressed mAbs were instead incubated at described conditions, in particular 0.2 – 20 μg/mL for anti-SARS-CoV-2 mAbs and 5 μg/mL for the rest of datasets. ELISA plates were washed three times with PBS. 50µL of secondary antibody (Jackson anti-mouse Fc HRP Cat. No. 115-035-008 or anti-human Fc HRP Cat. No. 109-035-008) were added and incubated for 1h at room temperature. The plates were then washed five times with PBS Tween 0.1%. The reaction was developed with 50µL of TMB (ThermoFisher 1-Step™ TMB ELISA Substrate Solution Turbo, Cat. No. 34022) at room temperature for 5 minutes and then neutralized with 50µL of 0.5M sulfuric acid. Finally, the absorbance was read at 450 nm using the EnVision plate reader.

### Affinity measurements by Biolayer Interferometry (Octet)

Antibodies to test were diluted at the concentration of 5 μg/mL in Octet Buffer (PBS, 0.02% Tween-20, 0.1% BSA). Antigens were used at the concentration of 100 nM in the same buffer. Test was performed in 384w titled well propylene microplates (Sartorius Cat. No. 18-5166) with 50 – 80 microliters of sample. Biosensors used were either AMC (anti-mouse capture, Sartorius, Cat. No. 18-5089) for mouse antibodies or AHC (anti-human capture Cat. No. 18-5060) for human antibodies. Assays were performed using Octet RED384.

### Graphs

Graphs were made using GraphPad Prism 9 and Adobe Illustrator. Alternatively, graphs were made in R and saved as SVG files, then imported in Illustrator.

### Gene Expression Heatmaps

GEX heatmaps were generated using the pheatmap R package. Data were retrieved from Seurat objects using the “data” slot, which contains log-normalized expression values. In pheatmap, average gene expression values were scaled across genes, with the data centered to have a mean of 0 and a standard deviation of 1.

### Statistics

Statistics were calculated in GraphPad, R or Excel. Populations were tested for normality using Kolmogorov–Smirnov test. If normally distributed, means were compared using a parametric test, Student’s t-test for homoscedastic samples and Welch’s t-test for heteroscedastic samples. When parametric, ANOVA one-way was applied for comparing multiple distribution, followed by a Dunnett’s test. Most of populations had a non-normal distribution, thus mostly non-parametric tests were used. Mann-Whitney U-test was applied to most experiments. Alternatively, a Kruskal-Wallis test followed by post-hoc multiple comparisons using Dunn’s test. P-values were typically corrected using the Benjamini-Hochberg FDR correction. To determine whether two distributions with categorical data (HIGH, MID, LOW) were statistically different, a Chi-squared test for independence was used (GraphPad). When samples were particularly small and belonging to only one dataset (OVA BC181), G-test was used in Rwith the function *DescTools::GTest().* In this case, Chi-squared test results were also reported in the figure legend.

### Protein antigens

All proteins were purchased from vendors except ST2 and FN14, which were produced in house (see **Recombinant Protein Production** section). Ovalbumin = InvivoGen OVA EndoFit Cat. No. vac-pova-100; Haemagglutinin (HA) = Influenza A H1N1 HA (A/Puerto Rico/8/1934) His-tag Recombinant Protein, ThermoFisher Scientific, Cat. No. A42599; ORF8 = SARS-CoV-2 ORF8 Protein His-tag, ACROBiosystems Cat. No. OR8-C52H1-100UG; RBD = SARS-CoV-2 (COVID-19) S protein RBD, His Tag, Cat. No. SPD-C52H3-100UG, ACROBiosystems; SPIKE COVID19 = SARS-CoV-2 (2019-nCoV) Spike S1+S2 ECD-His Recombinant Protein (100 ug), SinoBiological Cat. No. 40589-V08B1; Human coronavirus HKU1 (isolate N5) (HCoV-HKU1) Spike Protein, ACROBioSystems, Cat. No. 40606-V08B-SIB-100ug; Human coronavirus (HCoV-OC43) Spike Protein (S1+S2 ECD, His Tag), ACROBiosystems Cat. No. 40607-V08B-SIB-100ug; Human coronavirus (HCoV-NL63) Spike Protein (S1+S2 ECD, His Tag), Cat. No. 40604-V08B-SIB-100ug; Human coronavirus (HCoV-229E) Spike Protein (S1+S2 ECD, His Tag), Cat. No. 40605-V08B-SIB-100ug.

### Recombinant Protein Production

Protocols to produce ST2, FN14 and antibodies follow.

### Recombinant human ST2 extracellular domain

The sequence corresponding to human ST2 amino acids 19-328 (NCBI Reference Sequence: NP_003847.2) was cloned into pDEST12.2 via restriction sites using SbfI-HF (Cat. No. R3642L) and NheI-HF (Cat. No. R3131L) enzymes (NEB, New England Biolabs) and corresponding overhangs. The cloning reaction was performed using a Quick Ligation™ Kit (NEB, Cat. No. M2200L). An N-terminal CD33 signal peptide (MPLLLLLPLLWAGALA) was added to the ST2 construct as well as a C-terminal AviTag™ plus His10 tag (GGSGGSGGSGGSGLNDIFEAQKIEWHEAAHHHHHHHHHH) for expression, purification and biotinylation. Cloning success was then confirmed by sequencing (Source Bioscience). Recombinant ST2 protein expression was carried out in suspension-adapted CHO-K1 cells modified to express Epstein-Barr virus (EBV) nuclear antigen-1 (EBNA-1) provided by our internal cell bank. These cells were transiently transfected using a polyethylenimine (PEI) based protocol (R&D Systems PEI Star transfection reagent, Cat. No. 7854). For cell-line development details and transfection methods see reference^73^. Recombinant protein was then purified from filtered supernatant by Ni-affinity chromatography (HisTrap™ Excel 5 mL, Cytiva, Cat. No. 17371206) in 2xDPBS and eluted in 2xDPBS, 500 mM Imidazole pH7.2. Ni-affinity elution was then purified by size exclusion chromatography (HiLoad® 16/600 Superdex® 200 PG, Cytiva, Cat. No. 28989335) in 2x DPBS. Biotinylation of the AviTag™ was achieved via the addition of BirA using a commercial kit (Avidity) following manufacturer’s instructions. An additional size exclusion chromatography was performed, as previous, on biotinylated ST2 into 2xDPBS. Protein Mw, purity >95% and biotinylation were confirmed via SDS-PAGE and LC-MS. Protein concentration was calculated from absorbance at 280nm using extinction coefficient and Mw values from ProtParam – Expasy (https://web.expasy.org/protparam/). All proteins were flash frozen in liquid nitrogen and stored at -70°C.

### Recombinant human FN14

The sequence corresponding to Fn14 protein amino acids 29-80 (NCBI reference sequence NP_057723.1) was cloned into pEDST12.2. An N-terminal CD33 signal peptide (MPLLLLLPLLWAGALA) was added on the N-terminus as well as C-terminal Avi Tag and His10 tag. Recombinant FN14 protein was expressed in Expi293, with transient transfection using polyethylenimine (PEI, R&D Systems PEI Star transfection reagent, Cat. No. 7854). Cells were fed on day 0 and day 3, harvested on day 5. Recombinant protein was then purified from filtered supernatant by Ni-affinity chromatography (HisTrap™ Excel 5 mL, Cytiva, Cat. No. 17371206) in 2xDPBS and eluted in 2xDPBS, 500 mM Imidazole pH7.2. Ni-affinity elution was then purified by size exclusion chromatography (HiLoad® 16/600 Superdex® 75 PG, Cytiva, Cat. No. 28989333) in 2x DPBS. Biotinylation of the AviTag™ was achieved via the addition of BirA using a commercial kit (Avidity) following manufacturer’s instructions. An additional size exclusion chromatography was performed, as previous, on biotinylated FN14. Samples were concentrated to around 1mg/L, aliquoted, flash-frozen, and stored at -70C.

### Antibody synthesis, expression and purification

All proteins were expressed in suspension-adapted EB-GS22 CHO cells^73^ and maintained in AZ proprietary medium supplemented with methionine sulfoximine and hygromycin. Cells, growing in deep 24-well blocks, were transfected at a density of 4x10^6^/ml using PEI Max® (Polysciences Inc – Cat. No. 24765) at a PEI:DNA ratio of 5:1. Several hours post transfection, cells were temperature shifted to 34°C and shaken at 240 rpm in 5% CO2, 80% humidity for ten days. The conditioned media, containing secreted protein, was harvested on day ten by centrifugation at 2600 rpm for 30 minutes. The clarified media was purified using PhyNexus ProPlus LX PhyTips (Biotage – Cat. No. PTH-91-16-01) and buffer exchanged into PBS following elution using Zeba™ Spin Desalting Plates (ThermoFisher Scientific – Cat. No. 89808).

### Cell treatment with CB-6644, RNA extraction, cDNA preparation and qRT-PCR

1.5 million SP2 and RAMOS B-cell lines were treated with CB-6644 (Cayman Chemical Cat. No. CAY36125-10 mg) or DMSO for the specified amount of time and concentration. Cell pellets were then frozen at -70C until use. RNA was extracted using Qiagen RNeasy mini kit (Cat. No. 74104) according to manufacturer’s instructions. Briefly, pellets were resuspended in 350 microliters of Lysis Buffer, following 1 volume of 70% ethanol. Lysates were loaded onto provided columns and spun for 15s at 10,000 x g. After 1 volume wash with Buffer RW1, columns are treated with Qiagen RNase-Free DNase Set (Cat. No. 79254) for 15 min at RT. Afterwards, columns were washed 1x with 1 volume of RW1 buffer, following 2x RPE buffer washes (1x 15 s and 1x 2 min). Columns were dried by spinning 1 min at 10,000 x g in a clean collection tube. Finally, RNA was eluted in Nuclease-Free Water (NFW) and quantified at Qubit using RNA High Sensitivity Kit (ThermoFisher Scientific Cat. No. Q32852). 400 ng RNA per sample were retrotranscribed using High-Capacity RNA-to-cDNA™ Kit by ThermoFisher Scientific (Cat. No. 4387406) in a volume of 20 microliters. After RT, samples were diluted with NFW to 10 ng / microliter and used 1 microliter per well. qPCR was performed with PowerUp™ SYBR™ Green Master Mix for qPCR by Applied Biosystems (Cat. No. A25742) using the following amplification program: 50C hold 2 min, 95C hold 2 min; cycle 40x: 95C hold 1 s, 60 hold 30 s (Fast). Melting temperature curves were also generated to verify primer specificity. Primers used in this study were purchased from Integrated DNA Technologies (IDT), and relative Cat. No. are listed in the **Materials Used in This Study** Excel sheet.

## Supporting information

List of materials used in this study

## Authors Contributions

D.J.S. and X.R.R. originally ideated the project, supervised work, wrote the manuscript and provided funding for the project. D.A. supervised work and wrote the manuscript. M.C. conceptualised the study and the experiments, coordinated the team, performed all experiments except protein purification and immunisations, analysed data, performed all bioinformatic analyses, created figures, and wrote the manuscript. M.R. and S.G. produced recombinant proteins. R.M. and M.S. analysed bioinformatic data. V.C. performed experiments and analysed data. S.C. provided funding and supervised work.

## Acknowledgments

Sincere thanks to Matthew Burrell, Maximillian Dalglish and the BETXpress team of Biologics Engineering, AZ, for providing exceptional recombinant antibodies for this study. Thanks to the Animal Science & Technologies (AST) team for excellent *in vivo* work. Thanks to Marcin Wolny and Catherine Huntington for providing plasmids and proteins. Thank you, Franco Ferraro, for support provided with Octet. Thanks to Ozge Gizlenci for manuscript reading and comments. Thanks to Radhika Patel, Marjana Begum, Lilja Hardardottir and Raffaello Cimbro for the work done with sorting at AZ. Thanks to Andrew Buchanan, Luca Melchiori, Zenon Zenonos, Ziqi Long, Ben Dugan, Maria Tsoumpeli, Nachiket Shembekar, James Morris, Colin Hardman, Toby Gurran, Ziqi Zhou, Francesca Gasparrini, Rebecca Croasdale-Wood, DS’ team and XRR’s team for all the support and insightful discussion about this work. D.A. is supported by grants from the European Research Council (ERC-StG, B16 DOMINANCE, grant no. 850638); the Swedish Research Council (grant no. 2021-01164, 17 2021-01165) and the Knut and Alice Wallenberg Foundation (grant no 2021.0033 to D.A.)

---

**Supplementary Figure 1.**
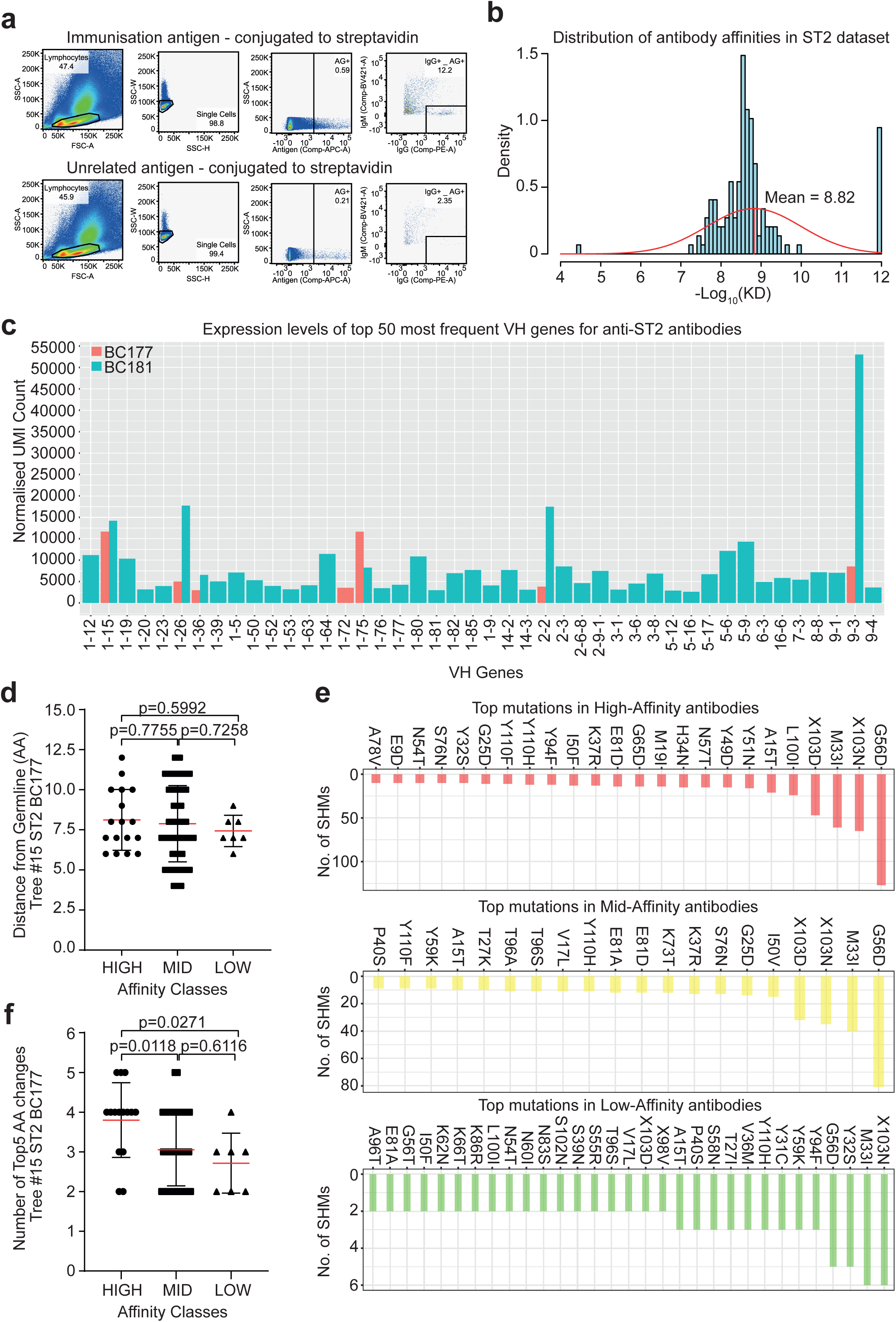
ST2 affinity datasets and SHM analyses. (a) Full sorting scheme for antigen-positive B-cells. Splenocytes or lymph node cells are gated on the most viable population, then from a single-cell gate antigen+/IgG+/IgM-cells are sorted. Gate for sorting is set on the same cells stained with unrelated antigen:streptavidin complex. (b) Affinity distribution in ST2 BC177 dataset. Most abundant affinity class (MID) was found to be nanomolar affinity (KD ∼ 10^-9^ M range or -Log_10_KD between 8 and 9). (c) Normalised UMI count of anti-ST2 antibodies V-genes. Comparison between ST2 BC177 and BC181 shows minimal overlap. V-gene 1-15 is among the most represented. (d) SHMs levels against affinity classes, measured as number of amino acid mutations accumulated with respect to germline in tree #15 (V-gene 1-15). Mann-Whitney U-test, two-tailed. N HIGH = 17, N MID = 68, N LOW = 7. Mean with SD is shown. (e) Frequence of common SHMs amino acid changes across affinity classes. (f) Number of top5 amino acid changes from (e) quantified, resulting in significant difference between HIGH/MID and HIGH/LOW. Mann-Whitney U-test, two-tailed. N HIGH = 15, N MID = 76, N LOW = 7. Mean with SD is shown.

**Supplementary Figure 2.**
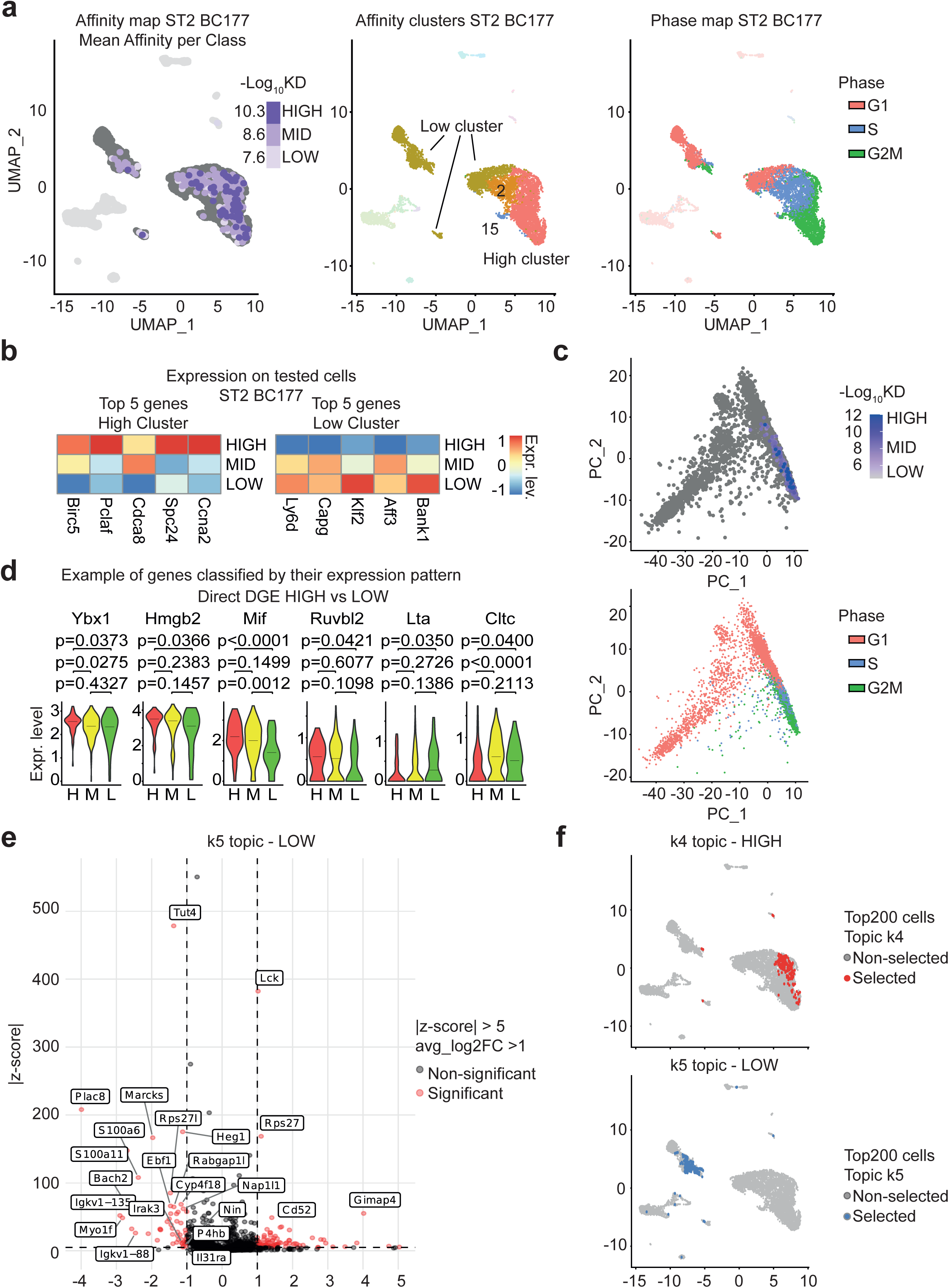
GEX data analyses. (a) Full affinity map of ST2 BC177 (left), division of B-cells in two affinity clusters, High cluster and Low cluster (centre), and segregation of cell cycle phases in analysed B-cells (right). (b) Verification of GEX trends found by DGE between High cluster and Low clusters in B-cells tested for affinity. (c) PCA plot of measured affinity shows clustering along a single principal component and shows better the homogeneous distribution of MID antibodies. (d) Markers selected based on their GEX characteristics derived from DGE analysis performed between HIGH and LOW cells. Mann-Whitney U-test, two-tailed. N HIGH = 58, N MID = 177, N LOW = 42. Black line indicates the median. (e) Volcano plot from Topic Model Analysis, topic k5. This topic is associated to lower affinity B-cells. (f) Top 200 cells involved in topic k4 and k5. HIGH topic (k4) involved mainly cells from GC DZ, while in topic k5 highest contributors were BMEM.

**Supplementary Figure 3.**
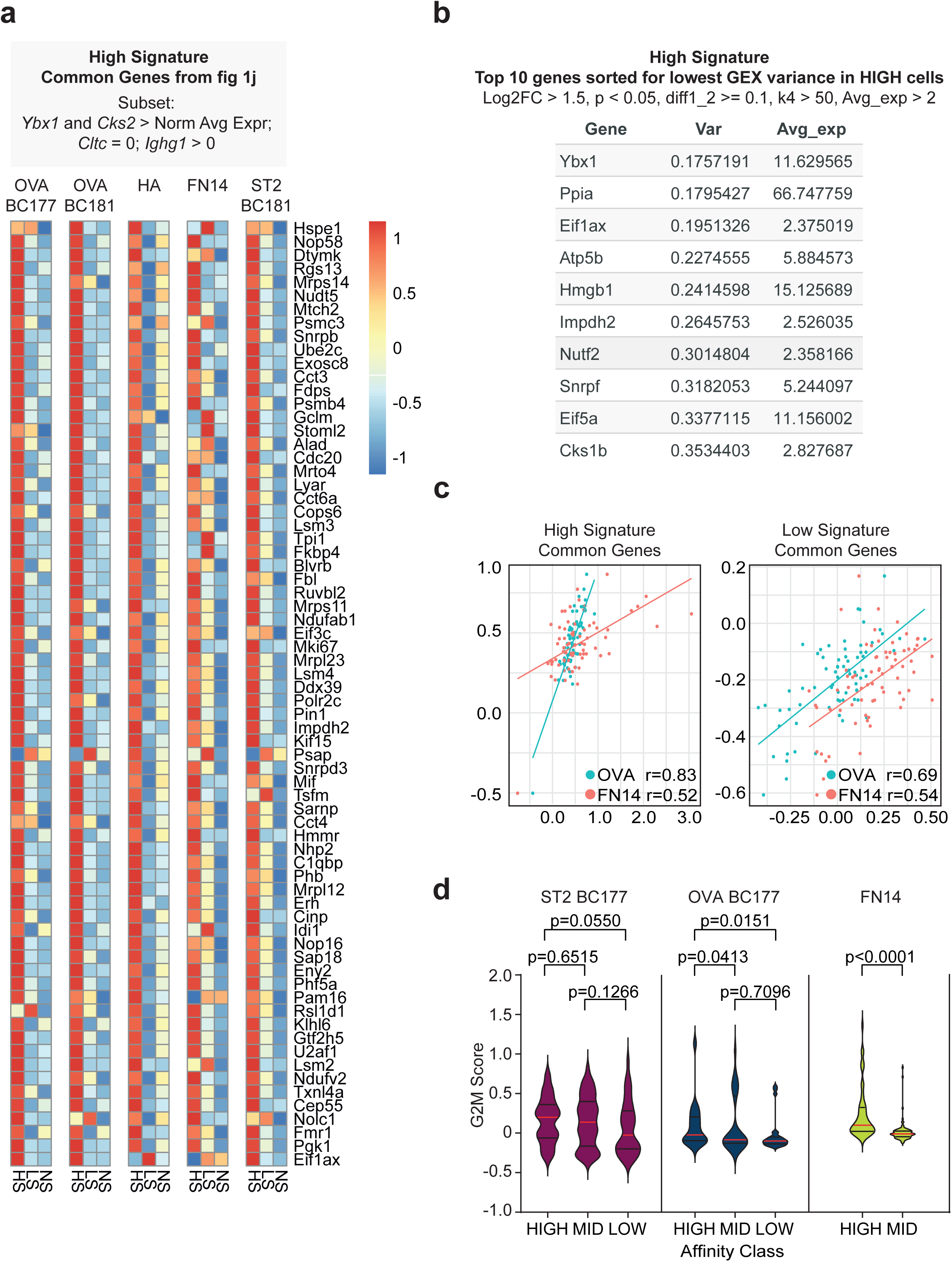
*High Signature* across datasets. (a) Heatmap of the 70 common genes found with all methods (Fig. 2j), across datasets. HS = *High Signature*, LS = *Low Signature*, NS = No Signature. (b) Table representing HS genes with lowest variance in high-affinity B-cells. (c) Correlation of all HS common genes across datasets. (d) Grouped violin plots showing G2M score in affinity-tested B-cells. G2M score was calculated using *CellCycleScoring* function in Seurat. Mann-Whitney U-test, two-tailed. ST2: N HIGH = 58, N MID = 177, N LOW = 42. OVA: N HIGH = 27, N MID = 36, N LOW = 32. FN14: N HIGH = 52, N MID = 97. Median with interquartile range is shown.

**Supplementary Figure 4.**
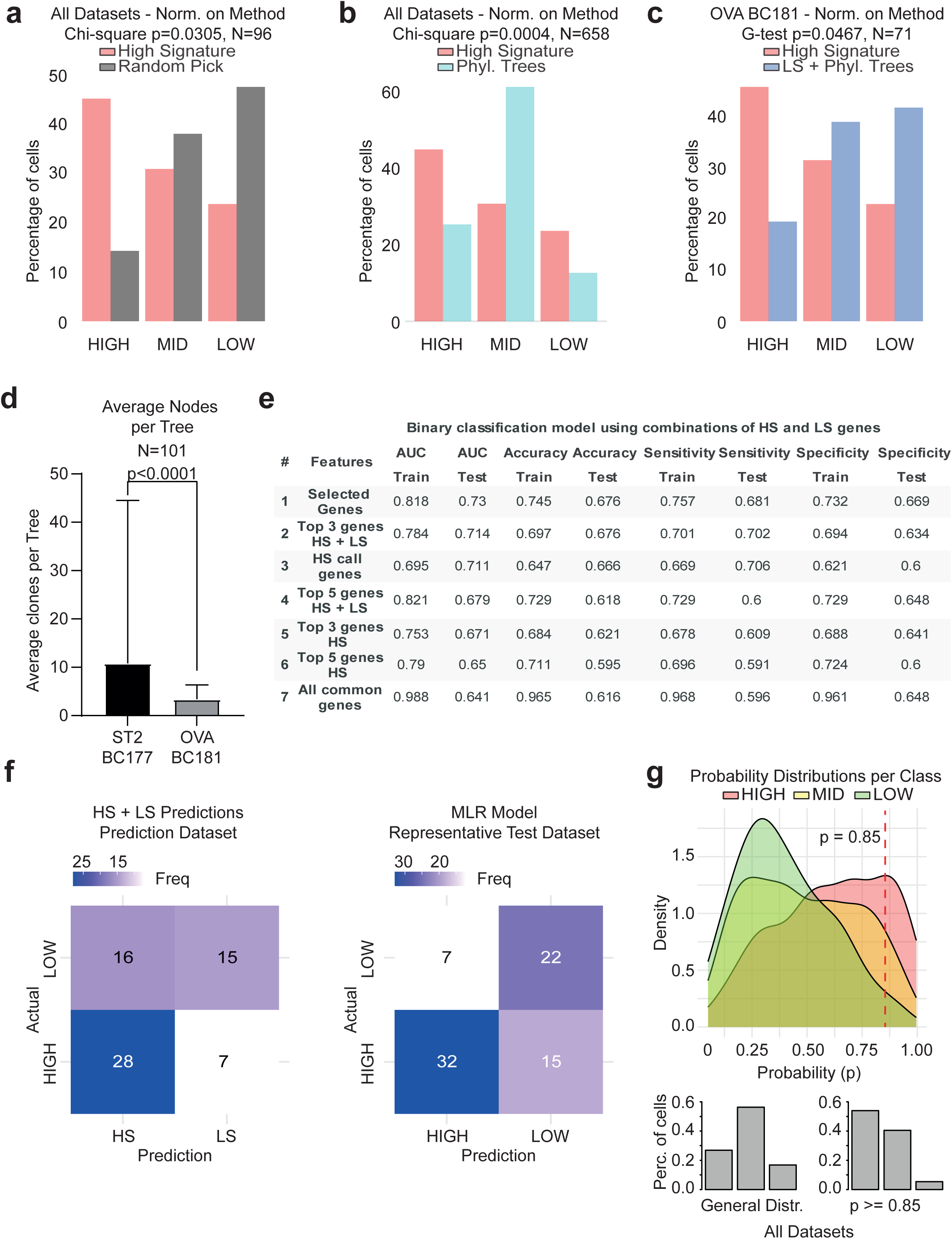
Predictions using HS and MLR model. (a-c) Comparison of different methods with HS. HS vs. Random Pick. N High Signature = HIGH 28, MID 26, LOW 16. N Random Pick HIGH 4, MID 10, LOW 12. Chi-squared test. (a). (b) Distribution of HS vs. phylogenetic trees from all datasets. Chi-squared test. N HS = HIGH 28, MID 26, LOW 16. N Phyl. Trees = HIGH 150, MID 363, LOW 75. (c) HS vs aggregated results obtained with LS + phylogenetic trees in OVA BC181 dataset. G-test shown. Chi-squared p-value = 0.0498. N HS = HIGH 16, MID 11, LOW 8. N LS + Phyl. Trees = HIGH 7, MID 14, LOW 15. (d) Average number of tree elements in ST2 BC177 dataset vs. OVA BC181. Mann-Whitney U-test, two-tailed. N ST2 = 55, N OVA = 46. Mean with SD is shown. (e) Optimisation of the machine learning model. Different subsets of genes deriving from the common genes or from top markers per method (Fig 2j) were used. Best results were obtained using genes listed in Figure 4i (Selected Genes). Total dataset N = 591. N HIGH = 158, N MID = 332, N LOW = 99. (f) Confusion matrixes for HS+LS predictions vs. MLR model. Data from Fig. 4d and from the Test Dataset of the selected MLR model. (g) Probability distributions per affinity class. Probabilities using selected binary MLR model were applied to all available datasets (including ST2) and classes. Density curves highlight separation across classes. Histograms in the bottom panel show how population is reshaped selecting cells with p >= 0.85, with 1 being HIGH.

**Supplementary Figure 5.**
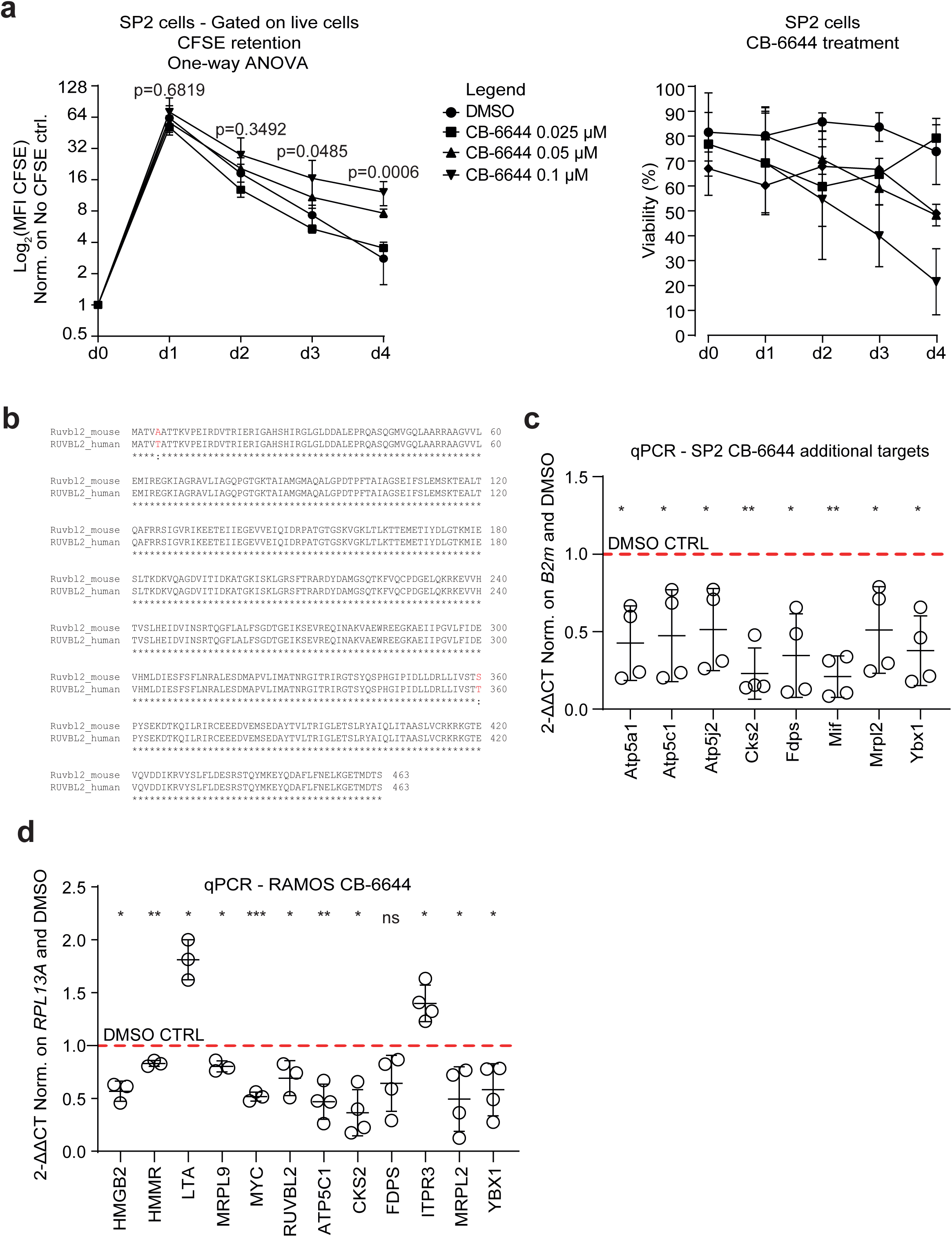
Inhibition of RUVBL2 protein in mouse and human B-cell lines. (a) CFSE retention titration curves showing difference in cell proliferation with DMSO occurs from day 2 post treatment (left graph) and Viability curves associated to different CB-6644 (RUVBL2 inhibitor) concentrations, measured by gating on most viable cells (right graph). One-way ANOVA. N DMSO = 5, N CB-6644 25 nM = 2, N CB-6644 50 nM = 2, N CB6644 100 nM = 3. For each point (mean), SD is shown. (b) Alignment of mouse Ruvbl2 and human RUVBL2 proteins showing 99.6% identity. (c) Additional targets measured by qPCR on SP2 B-cells. Paired t-test, two-tailed. 2^-ΔΔCT^ normalised on *B2m* and DMSO. Mean with SD is shown. (d) RT-qPCR on HS genes using RAMOS cells (human B-cell lymphoma) treated with 250 nM CB-6644 for 24h. Data show 2^-ΔΔCT^ normalised on *RPL13A* housekeeping gene and DMSO control (red dashed line = 1). Paired t-test, two-tailed. Mean with SD is shown.

**Supplementary Figure 6.**
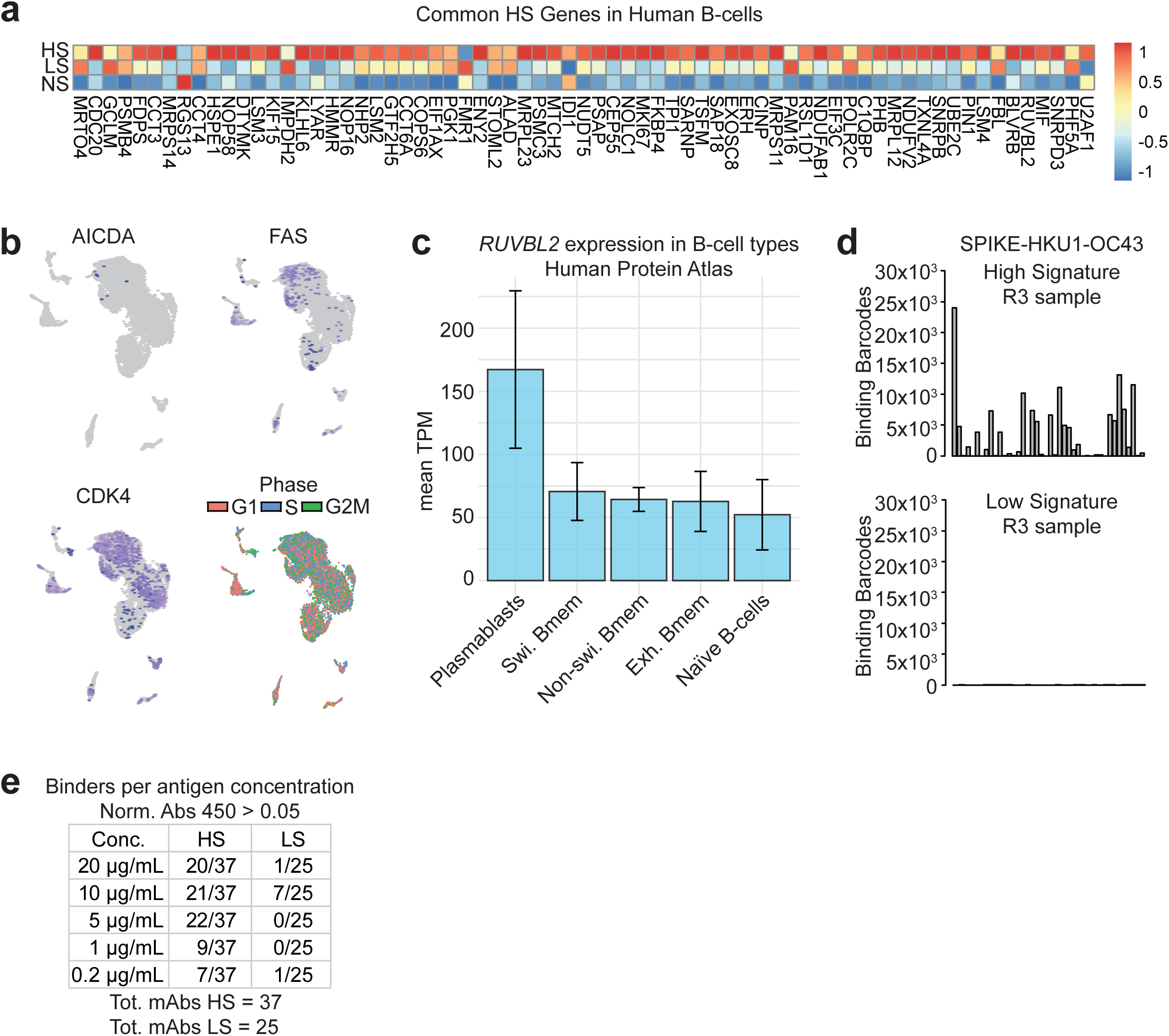
Human data. (a) Presence of HS in the re-analysed COVID human dataset^18^. HS = *High Signature*, LS = *Low Signature*, NS = No Signature. (b) Gene features of the COVID datasets (peripheral B-cells) show no segregation based on cell cycle phase and limited amount of GC activation markers as expected. (c) Human protein atlas data on RUVBL2 in several B-cell types. Swi. = Switched, Exh. = Exhausted. Mean with SD is shown. (d) Comparison of BCR-bound antigen:barcodes using HS vs LS calls in a specific patient (R3 sample). (e) Table summarising the number of antibody binders per given concentration. Data cut-off to define a binder was normalised absorbance 450 nm > 0.05. Independent experiment with respect to single-concentration ELISA of Fig. 6f.

